# Dissecting embryonic origins of brain pericytes, the role of angiogenesis, and fibroblasts’ contributions

**DOI:** 10.64898/2026.03.30.715397

**Authors:** Cynthia U Adjekukor, Katrinka M Kocha, Peng Huang, Sarah J Childs

## Abstract

Pericytes are mural cells that provide support to the endothelium of small blood vessels. Pericyte soma are regularly spaced along vessels, and their processes overlap only slightly. Given that vessel patterning is imprecise, we explore the interplay between vessel growth and pericyte recruitment that leads to even pericyte spacing. After recruitment to the zebrafish brain central arteries (CtAs), pericytes undergo rapid expansion, followed by morphological differentiation. Blocking angiogenesis by reducing Gpr124 (Wnt) or Vegf signaling reduces the length of the vessel network and the number of pericytes, preserving spacing, suggesting proportional recruitment of pericytes to cover the network and the territorial nature of pericytes. However, these initial brain pericytes have low proliferation rates. We demonstrate that additional pericytes are recruited firstly through migration of *col5a1-* and later *col1a2*-expressing fibroblasts into the brain. These second-wave pericytes retain some fibroblast properties and show elevated *col1a2* levels in a model of pericyte loss (*notch3* mutants). Our data provide new insights into the developmental timing, expansion, and novel origins of late-arriving brain pericytes during embryogenesis.

**SUMMARY STATEMENT:** This article demonstrates that brain pericytes originate from multiple sources, including fibroblast-derived populations, and how pericyte numbers are adjusted in proportion to vessel development.

## INTRODUCTION

The revolution in single-cell sequencing has highlighted the diversity of cells in the brain, especially of vascular mural cells, which include pericytes, vascular smooth muscle cells, and other perivascular cells, such as fibroblasts, astrocytes, and immune cells (Yang et al., 2022). However, knowledge of their developmental lineages and relationships is sparse. Brain pericytes wrap around the endothelial lining of small blood vessels to stabilize them, regulate blood flow, and maintain blood-brain barrier integrity (Brown et al., 2019). They cover about 80 – 90% of the adult brain capillaries, and defects in their coverage have been implicated in intracranial hemorrhage, cerebral small vessel disease (SVD), and stroke (Armulik et al., 2010; Berthiaume et al., 2022; Gautam and Yao, 2018; Sun et al., 2021).

Pericytes and endothelial cell development are coupled. While endothelial cells are formed first through vasculogenesis and angiogenesis, they interact with Pdgfrβ+ve pericytes via secreting Pdgf-B, inducing migration, differentiation, and proliferation of pericytes on the growing vascular network (Benjamin et al., 1998; Hellström et al., 1999; Lindahl et al., 1997). Pericytes are characterized morphologically by having a cell body (soma) and typically two cytoplasmic processes that run along the endothelium (Armulik et al., 2011; Grant et al., 2019). These processes elaborate further over development depending on the size and type of vessel (Berthiaume et al., 2018; Grant et al., 2019). Pericytes migrate into the brain as *nkx3.1^+ve^*^;^ *pdgfr*β*^low^* precursors and differentiate to *pdgfr*β*^high^* cells that expand (Ahuja et al., 2024; Ando et al., 2019; Ando et al., 2021). It has been assumed that pericytes expand through proliferation. For instance, cultured mouse aortic rings suggest that proliferation is the primary mechanism by which pericytes expand their population. However, in this in vitro system, there is no other potential source of pericytes, in contrast to development (Chiaverina et al., 2019). In the zebrafish trunk, mural cells on the dorsal aorta also increase by proliferation (Ando et al., 2016).

During angiogenesis, pericytes become activated, extending their process to move in the direction of the sprouting vessel (Dessalles et al., 2021; Payne et al., 2021). They are highly territorial, extending their processes along the vessels with either no or very small overlap with those of neighboring pericytes (Berthiaume et al., 2018; Graff et al., 2025). These processes migrate along the endothelial junction, making direct contact with it throughout embryogenesis (Ando et al., 2016). Both adult and aged brains show comparable levels of brain pericyte coverage in the mouse cortex, suggesting that pericyte coverage and spacing might be established during development (Berthiaume et al., 2022).

Brain pericyte precursors or intermediate progenitors originate from the cranial neural crest and mesoderm in mammals and zebrafish (Ahuja et al., 2024; Ando et al., 2016; Etchevers et al., 2001; Pouget et al., 2008). We and others have shown that pericyte precursor development is regulated by the transcription factor *nkx3.1*, in both neural crest and mesoderm (Ahuja et al., 2024). Since pericytes can differentiate from various lineages, pericyte precursors are clearly diverse. To further diversify the population, single-cell sequencing data from pericyte precursors showed that intermediate pericyte progenitors express fibroblast markers. Like pericytes, meninges are also derived from mesoderm and neural crest, and they contain many resident fibroblasts (Jiang et al., 2002). In mice, meningeal fibroblasts enter the brain along vessels and take up perivascular locations (Jones et al., 2023). Some of these perivascular fibroblasts are double positive for Collagen-1 and *PDGFR-*β (Jones et al., 2026).

Fibroblasts also act as progenitors of pericytes in the developing embryonic mouse skin and intersegmental vessels (ISVs) of zebrafish (Goss et al., 2021; Rajan et al., 2020). Collagen expression is a characteristic of fibroblasts, and mouse brain fibroblasts express common fibrillar collagens, *Col1a1*, *Col1a2*, and *Col5a1* genes (Vanlandewijck et al., 2018), as do zebrafish trunk fibroblasts (Rajan et al., 2020). Single-cell RNA sequencing of prenatal human brains shows fibroblasts and pericytes both express *COL1A1* and *COL1A2* (Crouch et al., 2022; Speir et al., 2021). Differentiated pericytes from human cell *NKX3.1* precursors also express *COL1A1* and *COL1A2* (Lee et al., 2024). Single-cell RNA sequencing data from 5 dpf zebrafish pericyte-labelled (*pdgfr*β*+ve*) cells show *col1a2* expression in the pericyte cluster (Shih et al., 2021). Thus, it is evident that fibroblast markers are expressed in pericytes. However, few studies have examined whether fibroblasts give rise to brain pericytes.

Here, we investigate the developmental origins and trajectories of brain pericytes in zebrafish to gain a comprehensive understanding of their lineage. We study the stage at which pericytes reach a dense layer of vessel coverage, the role of endothelial signaling in this process, and the sources of pericytes. We demonstrate that the dense level of brain pericyte coverage achieved at the end of embryogenesis is similar to that of larval stages. We show that pericyte spacing is conserved in normal and disturbed angiogenesis. Additionally, we show that brain pericyte proliferation rates are low, indicating that continuous cell differentiation is required for pericyte expansion in post-embryonic stages. We identify a previously undescribed origin of brain pericytes via the migration of brain fibroblasts that is spatiotemporally regulated: *col5a1*-expressing fibroblasts give rise to early pericytes around the hindbrain midline, whereas later embryonic pericytes from the meninges derive from both *col5a1*- and *col1a2*- expressing fibroblasts. Finally, we examined the SVD *notch3^fh322^* mutants and found that their pericytes exhibit elevated *col1a2* mRNA expression, an early indicator of fibrosis.

## RESULTS

### A burst of pericyte expansion occurs during late embryonic development

Since pericytes almost completely cover capillaries in adulthood, we sought to understand how their population expands to cover the vessel network during angiogenesis from embryonic through larval stages using live-imaging of transgenic zebrafish with reporters for pericytes, *TgBAC(pdgfrb:Gal4FF)^ca42^; Tg(5xUAS:EGFP)^nkuasgfp1a^*, and endothelial cells, *Tg(kdrl:mCherry)^ci5^*. We imaged vessels of the mid- and hindbrain where angiogenic Central Arteries (CtAs) are located. These capillary-sized vessels are initially ‘naked’ but are progressively covered by pericytes over development. Differentiating pericytes show a rounded soma with processes extending along the endothelium in both mid- and hindbrain CtAs at 3 dpf (Fig. 1A-C). As the blood vessel network develops slightly earlier than pericyte emergence, we assessed two metrics: pericyte coverage (the total length of pericyte soma plus processes over the total vessel network length) and pericyte density (the number of pericytes per vessel network length) (Fig. 1D). Total vessel network length was calculated using Vessel Metrics (McGarry et al., 2024) and pericyte length was measured along the longitudinal axis of the vessel (Fig. S1A-C).

**Fig. 1.**
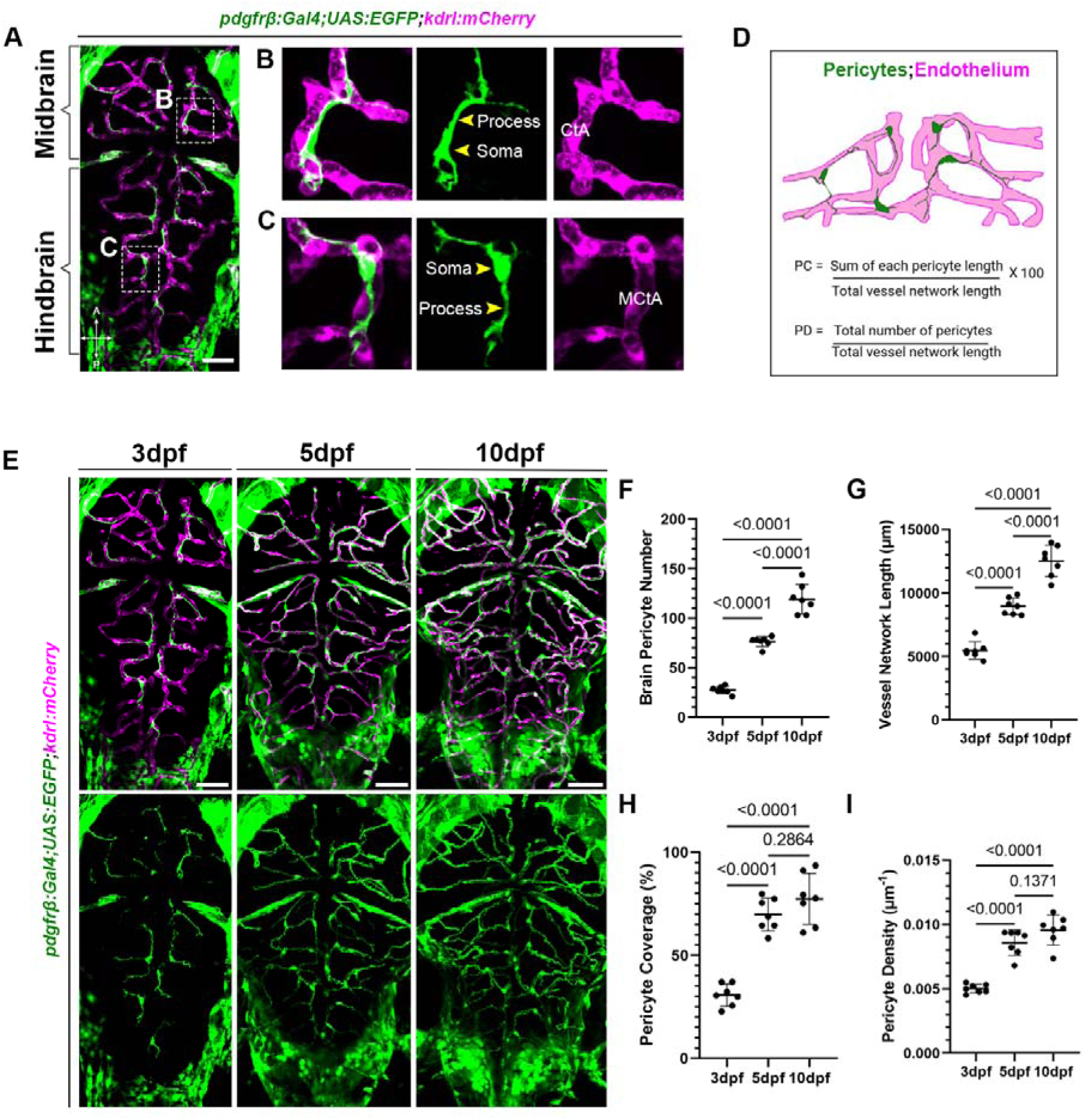
Identification of critical embryonic periods for pericyte recruitment and differentiation. (A) Dorsal view of 3dpf zebrafish larva on transgenic lines for pericytes (green) and endothelial cells (magenta) showing pericyte morphology and location in the small vessels of the (B) midbrain and (C) hindbrain. (D) Schematic of how pericyte coverage (PC) and pericyte density (PD) were measured. (E) Confocal images of a single zebrafish larva imaged repetitively at 3, 5, and 10dpf, showing pericyte and endothelial network development. (F-I) Quantification of (F) pericyte number, (G) vessel network length, (H) pericyte coverage, and (I) pericyte density of individual zebrafish larvae serially imaged at 3, 5, and 10 dpf (n= 7). CtA – central arteries, MCtA – mesencephalic central arteries. Statistical analysis was performed using a one-way ANOVA with Tukey’s multiple-comparison test, with results represented as mean ± sd. Scale bars: 50µm.

When we serially imaged the same embryos at 3, 5, and 10 dpf, we observed a progressive increase in brain pericyte number, from an average of 28 pericytes/brain at 3 dpf to 120 pericytes/brain at 10 dpf, representing an approximately 300% increase (Fig. 1E-F). Interestingly, both vessel network length and pericyte numbers appear to increase steadily from 3 through 10 dpf as the animal grows (Fig. 1G). In contrast, pericyte coverage and density on vessels are initially low, but reach a plateau between 5 and 10 dpf. Pericyte processes also elaborate over this period. At 3 dpf, a stage preceding blood-brain barrier formation, pericytes cover approximately 30% of the vessel network, with an average density of 0.005 µm**^-1^** (∼ 1 cell/200 µm of vessel length). By 5 dpf, pericyte processes extend to cover approximately 70% of endothelium, with an increased average density of 0.0086 µm^-1^ (∼1 cell/116 µm of vessel length). Neither pericyte coverage nor density changes significantly from these values at 10 dpf (Fig. 1H-I). Our data suggests that the period between 3 and 5 dpf is a time of rapid pericyte expansion. Since pericyte coverage in early development lags vessel growth, this period requires rapid pericyte recruitment and differentiation. After embryogenesis, pericytes and endothelium undergo more proportional expansion. Also, our data highlights the period between 3 dpf and 5 dpf as a critical time when pericyte numbers increase, and pericytes extend their processes to cover growing vessels. Thus, zebrafish brain pericytes achieve a relatively static vessel coverage ratio at 5dpf, coincidental with blood-brain barrier formation (O’Brown et al., 2019).

### Role of endothelium in the migration and differentiation of the initial pericyte pool

Next, we inquired about the relationship between pericyte spacing and endothelial network length. For instance, is there a fixed pool of pericyte progenitors waiting to populate the endothelium, or is there a dynamic adjustment of pericyte differentiation that responds to angiogenesis? To test this, we used inhibitors to reduce the length of the endothelial network. CNS angiogenesis is uniquely regulated by Wnt7a/b signaling via Gpr124 (Posokhova et al., 2015; Vanhollebeke et al., 2015; Zhou and Nathans, 2014). Wnt signaling is active in endothelial cells at 48 and 72 hpf, as shown by signal from Wnt reporter *top:GFP* colocalized with endothelial *kdrl:mCherry* in the CtAs (Fig. 2A-D). On the contrary, Wnt signaling is not activated in pericytes, as shown by the lack of co-expression of the *top:GFP* and *pdgfrb^NTR-mCherry^* reporters (Fig. 2E-F).

**Fig. 2.**
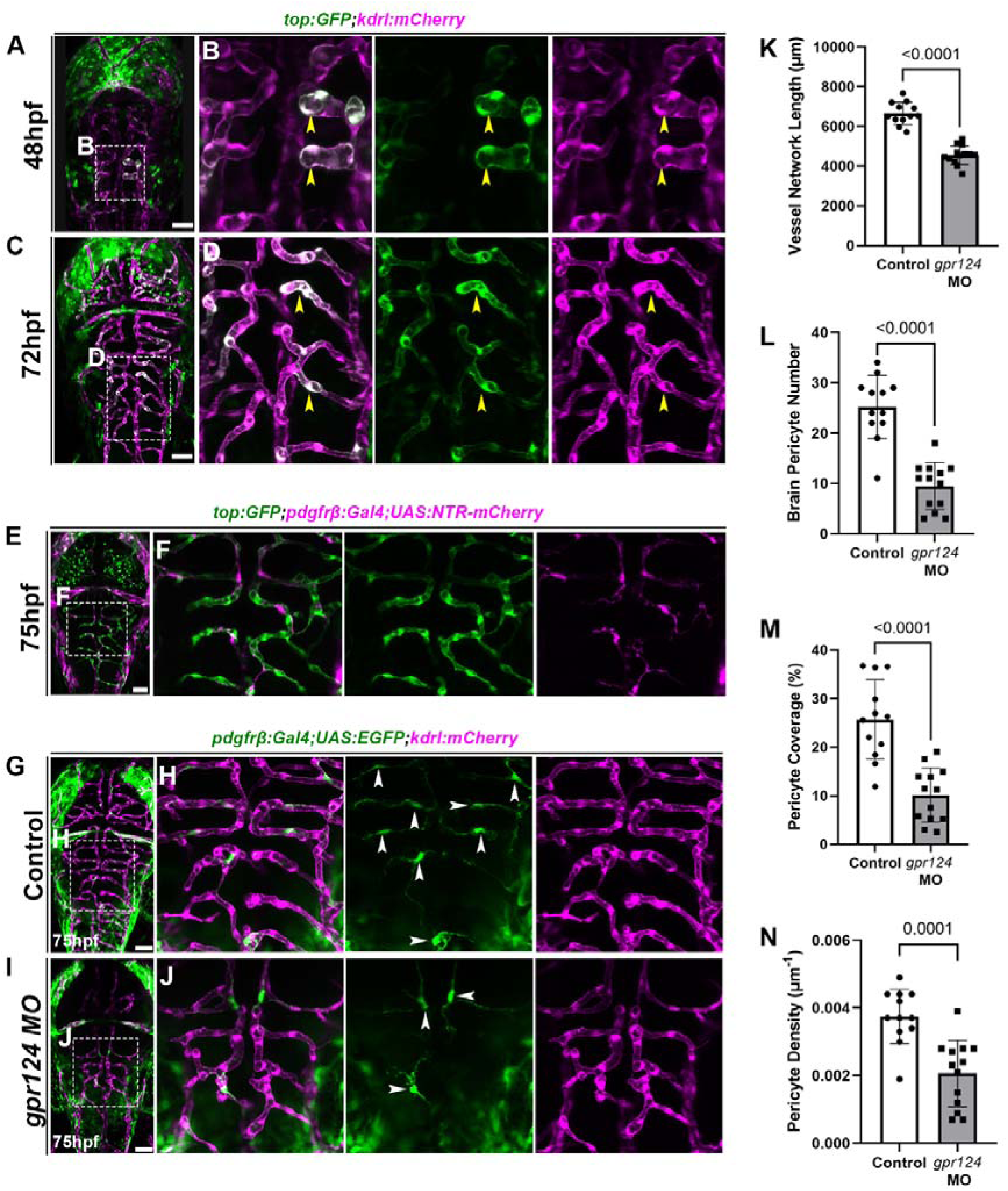
Blocking angiogenesis through reducing *gpr124* expression strongly reduces pericyte numbers on brain vessels. (A-D) Dorsal image of zebrafish larva at (A-B) 48hpf and (C-D) 72hpf on Wnt reporter (green) and endothelial transgenic (magenta) lines. Yellow arrows on the inset images indicate Wnt activation in the CtAs. (E) 75hpf larva on Wnt reporter (green) and pericyte (magenta) transgenic lines. (F) An inset of the hindbrain shows no colocalization of the Wnt reporter with pericytes. (G-J) Confocal images of 75hpf (G-H) control and (I-J) *gpr124* morpholino (MO) injected larvae on pericyte (green) and endothelial (magenta) transgenic lines. White arrows on the inset images point to pericytes on the hindbrain. (K-N) Quantification showing the *gpr124* MO groups had significantly reduced (K) vessel network length, (L) pericyte number, (M) pericyte coverage, and (N) pericyte density. (n: control= 12, *gpr124* MO= 13). All statistical analyses are unpaired t-tests represented as mean ± sd. Scale bars: 50µm.

To test the effect of Wnt signaling on pericyte development, we knocked down *gpr124* expression using a morpholino (Vanhollebeke et al., 2015). As expected, the uninjected controls and *gpr124* morphants (MO) show no morphological abnormalities at 75 hpf (Fig. S2A). However, *gpr124* morphants have fewer CtAs and lack Wnt activity in the endothelium, while reporter expression in non-vascular brain cells remains unchanged, confirming the specificity of the *gpr124* phenotype to the endothelium (Fig. S2B-C). At 75 hpf, the average vessel network length decreased by ∼ 32% in the *gpr124* MO group (avg: control= 6659.92, *gpr124* MO=4557.18, Fig. 2G-K). Additionally, *gpr124* MO embryos have fewer brain pericytes, with an average of 9 pericytes/brain, compared to 25 pericytes/brain in control embryos, a 64% reduction from wild-type numbers (Fig. 2L). Since the number of pericytes decreases with vessel network length, this suggests a dynamic adjustment in pericyte recruitment/differentiation in response to reduced angiogenesis. Pericyte coverage decreased by 60% and density by 42% in the *gpr124* MO group, suggesting the pericytes are insufficient to cover the vessels (Fig. 2M-N).

Furthermore, we independently tested the role of the endothelium by selectively inhibiting Vegfr1/2-mediated CtA sprouting using the inhibitor SU5416, starting at 32 hpf when CtA sprouting has just begun, and most large-caliber vessels (LDA, PHBC, MCev, MSV) are fully formed (Fig. S3A) to selectively inhibit brain CtA angiogenesis. Consistent with the *gpr124* knockdown model at 75hpf, SU5416-treated embryos exhibit decreased blood vessel network length, pericyte number, pericyte coverage, and density compared to the DMSO group (Fig. S3B-I).

Taken together, targeting two pathways that block CtA angiogenesis suggests an interdependence between pericyte recruitment/differentiation and angiogenesis. With reduced CtA angiogenesis, pericyte numbers and pericyte process extension on vessels are sharply reduced. However, pericytes were well spaced and maintained their territories on vessels.

### Brain pericytes proliferate to increase their numbers as the vessel network grows

We next examined the relationship between migration/differentiation and proliferation during pericyte population expansion. We used 25-hr time-lapse imaging of hindbrain CtA vessels in *pdgfrb^GFP^;kdrl:mCherry* embryos. The hindbrain CtAs are highly patterned, allowing tracking of pericyte migration and proliferation. We observed proliferative cells dividing, but, unexpectedly, far more cells ‘appeared’ on the CtA than proliferated. Beginning at 52 hpf, a time point before *pdgfrb^high^-*expressing pericytes are on the CtAs, and continuing to 77 hpf, 15.4% of pericytes on the CtAs divide. In contrast, 84.6% of pericytes appeared on the CtAs, reflecting cells differentiating to express *pdgfrb^high^* transgene while migrating into the brain from ventral sources (n= 5, avg hindbrain pericyte no= 7, Movie 1, Fig. 3A).

**Fig. 3.**
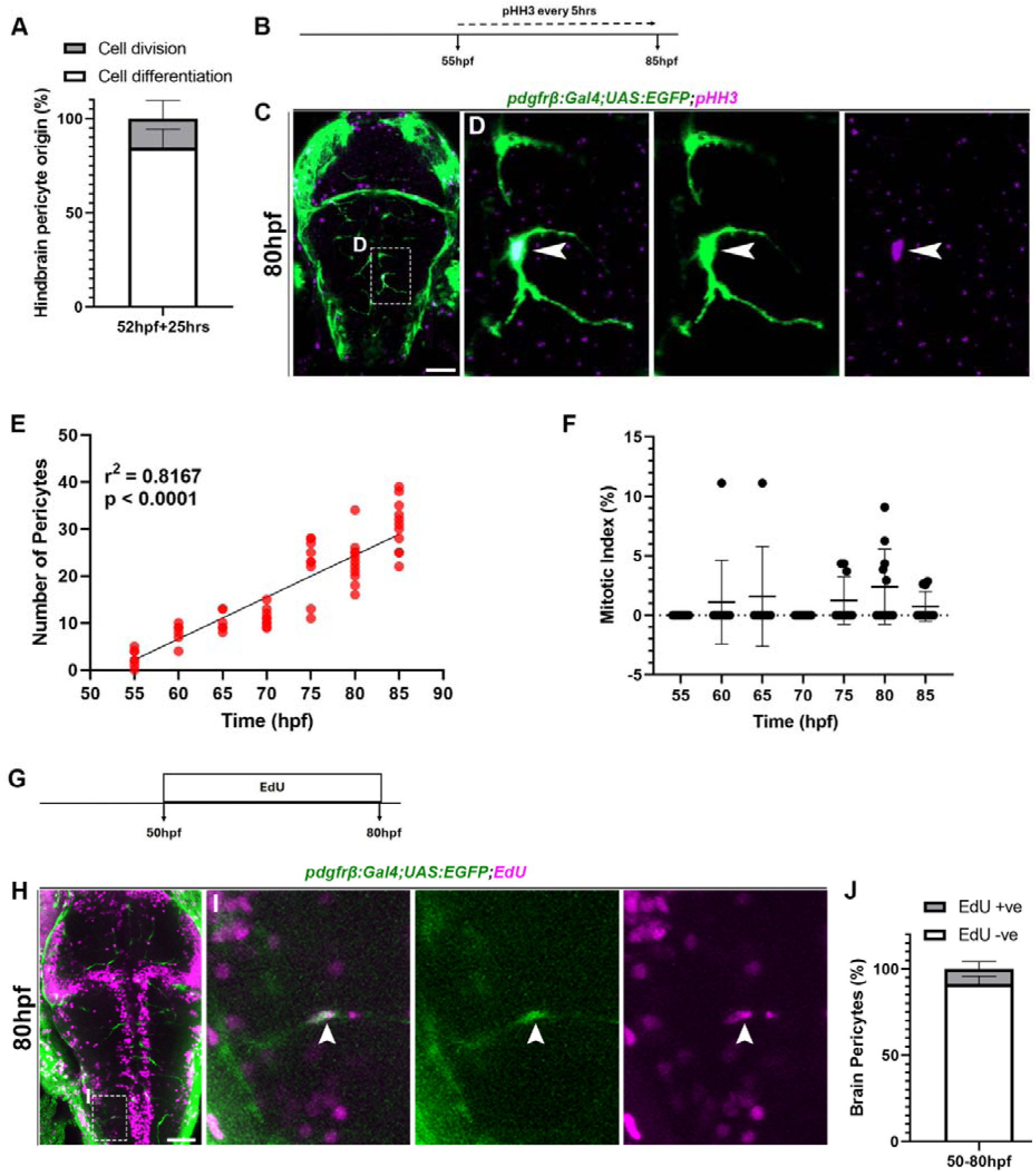
Proliferation contributes to increasing brain pericyte number. (A) Analysis of hindbrain pericyte proliferation vs. differentiation taken from a 25hr time-lapse movie shows the percentage of pericytes that divide (proliferate) or differentiate on the vessels. (B) Schematic of the sampling schedule for Phospho-histone H3 (pHH3) antibody staining (C-D) 80hpf larva on pericyte (green) line stained with pHH3 antibody. The white arrow shows co-expression of pericyte (green) with pHH3-positive nuclei (magenta), indicating a proliferating pericyte. (E) Linear regression analysis of pericyte number across the pHH3 staining intervals. (F) Graph of mitotic index defined as the average number of pHH3 positive pericytes per embryo across the time intervals (n: 55hpf= 11, 60hpf=10, 65hpf= 7, 70hpf= 11, 75hpf= 10, 80hpf= 11, 85hpf=11). (G) Schematic of EdU staining (H-I) 80hpf larva on pericyte (green) line stained with EdU (magenta). The white arrow indicates co-expression of pericyte soma and EdU staining. (J) Graph showing the percentage of brain pericytes positive or negative for EdU stain (n=20). Scale bars: 50µm.

We next assayed pericyte proliferation using the M-phase marker Phospho-histone H3 (pHH3) staining on *pdgfrb^GFP^* embryos. A time range of 55–85 hpf was sampled in 5 hr intervals (Fig. 3B). For most 5 hr intervals, at least one brain pericyte per embryo group is positive for pHH3, as shown at 80 hpf (Fig. 3C-D). Linear regression analysis showed pericyte numbers increase from 55- 85 hpf with a slope of 0.8899, and r^2^=0.82 (Fig. 3E). We calculated the mitotic index, defined as the percentage of pHH3-positive pericytes divided by the total number of pericytes per embryo at every stage. The highest mitotic index is at 80 hpf (2.41%), and the average mitotic index across all stages is 1.01% (Fig. 3F). Thus, the pericyte division rate is surprisingly low.

To independently test the proliferation rate, we incorporated the S-phase marker EdU into *pdgfrb^GFP^* embryos (Fig. 3G) over a 30-hr period from 50-80 hpf. We observe 8.6% of pericytes are positive for EdU within this longer timeframe (Fig. 3H-J). EdU incorporation supports the observation that proliferation is slow. Taken together with the observation of pericyte ‘appearance’ during time-lapse, this suggests that continuous differentiation and/or migration occurs during embryogenesis and accounts for the majority of ‘new’ pericytes.

### *col5a1-*expressing fibroblasts give rise to the early brain pericyte populations

What are the sources of new pericytes into the brain at later stages? These could arise from in situ differentiation of precursors or from significant in-migration of pericytes from elsewhere. Our recent single-cell sequencing analysis showed that pericyte intermediate progenitors at 30 hpf are closely related to fibroblasts and express the fibroblast markers *col1a2 and col5a1* (Ahuja et al., 2024). This data, along with data from DanioCell, shows that *col5a1* expression is similar to when pericyte progenitors arise (Sur et al., 2023). Therefore, we tested whether and when *col5a1*-expressing fibroblasts contribute to brain pericytes using transcriptional reporters *col5a1^NTR-mCherry^ for fibroblasts* and *kdrl:EGFP* for endothelium. At 32hpf, during the emergence of CtAs, *col5a1* fibroblasts were not attached to any hindbrain vessels nor the mid-cerebral vein (Fig. S4A-B). However, by 2dpf, we observe the emergence of *col5a1+ve* pericyte-like cells associated with the hindbrain CtAs, and by 3dpf, these cells have populated the hindbrain CtAs (Fig. 4A-D). A 25-hr time-lapse movie conducted from 48 hpf shows that *col5a1*-expressing pericyte-like cells migrate and divide along the hindbrain CtAs, similar to *pdgfrb*- and *nkx3.1*-expressing cells (Ahuja et al., 2024) (Movie 2). This consistency in timing, morphology, and pericyte-like behavior suggests that *col5a1* is likely expressed in immature pericyte progenitors that migrate to the hindbrain.

**Fig. 4.**
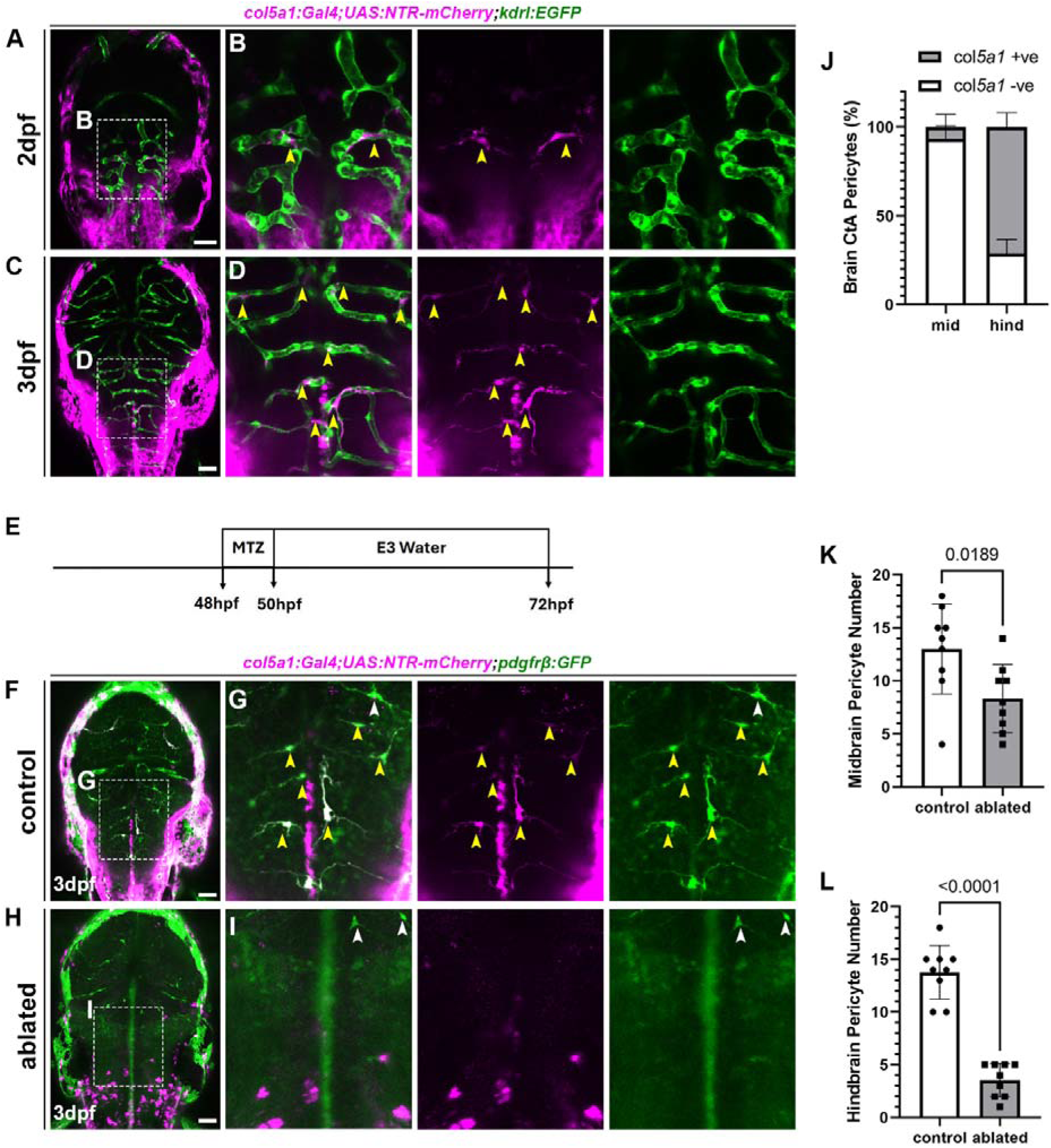
*col5a1*-expressing cells differentiate into early embryonic brain pericytes. (A-D) Dorsal view of a (A-B) 2dpf and (C-D) 3dpf larva on *col5a1* (magenta) and endothelial (*kdrl*; green) reporter lines. Yellow arrows indicate pericyte-like cells on the endothelium expressing *col5a1*. (E) Schematic of metronidazole (MTZ) drug treatment. (F-I) Dorsal view of a (F-G) control and (H-I) MTZ-ablated 3dpf larva *col5a1* and *pdgfr*β transgenic lines. Yellow arrows in the inset show *col5a1*+ve pericytes, while white arrows indicate *col5a1*-ve pericytes. (J) Graph showing the proportion of CtA pericytes that are positive or negative for *col5a1* expression at 3dpf (n=9). (K-L) Quantification of (K) midbrain and (L) hindbrain pericytes shows significantly reduced pericyte numbers after ablation of col5a1-expressing cells using MTZ (n: control & ablated = 9). Statistical analyses are unpaired t-tests represented as mean ± sd. Scale bars: 50µm.

To understand how critical *col5a1-expressing* cells are to brain pericyte development, we performed metronidazole ablation of *col5a1^NTR-mCherry^-*expressing cells in transgenic pericyte *pdgfrb;GFP* reporter from 48 hpf to 72 hpf (3 dpf) (Fig. 4E). In the control group, we observed co-expression of *col5a1* and *pdgfr*β reporters in the mid- and hindbrain CtAs at 3dpf, while no co-expression of markers was observed in the ablated group, suggesting that all double-positive cells of *col5a1* and *pdgfrb* were eliminated by the treatment (Fig. 4F-I). Interestingly, while hindbrain *col5a1*+ve pericytes are located near the midline of the brain, the midbrain *col5a1*+ve pericytes are lateral and proximate to the meninges. Specifically, about 7% of midbrain and 71% of hindbrain pericytes co-express the *col5a1* and *pdgfr*β markers in the control group at 3dpf (Fig. 4J). Furthermore, we observe a reduction in brain pericytes in both the mid- and hindbrain of the ablated group, with a greater reduction in hindbrain pericytes, suggesting *col5a1*-expressing pericytes are an important contributor to overall pericyte numbers, particularly in the hindbrain (Fig. 4K-L).

To test whether this early reduction in pericytes had a lasting phenotype, we performed similar metronidazole ablation in *col5a1;^NTR-mCherry^*and *pdgfr*β*;GFP*, and imaged them later. In wildtype animals, there is co-expression of both markers in the mid- and hindbrain CtAs at 5dpf (Fig. S4C-E). After ablation of *col5a1*-expressing cells using metronidazole treatment from 48-50hpf, we assayed pericytes at 5dpf (Fig. S4F-J). We observed that only about 3.87% of midbrain and 39.12% of hindbrain pericytes co-express the *col5a1* and *pdgfr*β markers in the control group at 5dpf (Fig. S4K). We also observed a significant reduction in hindbrain pericytes in the ablated group, with no change in the midbrain pericytes (Fig. S4L-M). This also supports that *col5a1*-expressing pericytes are important contributors of early migrating pericytes, especially in the hindbrain.

### Some late-emerging brain pericytes originate from a *col1a2*-expressing population

During zebrafish intersegmental vessel development, fibroblasts that are double-positive for *col5a1* and *col1a2* are pericyte progenitors (Rajan et al., 2020). Hence, we tested whether *col1a2*-expressing fibroblasts contribute to brain pericytes using a *col1a2* transcriptional reporter, *col1a2^NTR-mCherry^*, and *kdrl:EGFP* to mark endothelium. We observed large differences in the expression of the *col1a2* reporter compared with the *col5a1* reporter. At 32hpf which marks the beginning of brain angiogenesis, *col1a2*-expressing fibroblasts surround large vessels in the brain, the mid-cerebral (MceV) and mesencephalic veins (MsV) (Fig. S5A-B, Movie 3). This is interesting, as the *col5a1* reporter line showed no expression in these large vessels at this time. However, unlike the *col5a1-expressing cells*, *col1a2*-expressing pericyte-like cells are not observed on the hindbrain CtAs at 2dpf (Fig. S5C-D), but are present at 3dpf in the dorsal brain, a spatially distinct region. These cells show processes projecting into the brain, with similarities to pericyte processes connecting the meninges to midbrain CtAs (Fig. 5A-B). From serial images of the same embryo at 4dpf, we observe that the cell soma of the *col1a2^NTR-mCherry^* lineage cell moves away from the meninges and takes on a typical pericyte morphology, with a round soma with its process running longitudinally along the vessel (Fig. 5C-D). Time-lapse microscopy of *col1a2^NTR-mCherry^* lineage cells from 52 - 76hpf (Movies 4) shows double-positive *col1a2* and *pdgfr*β cell pericyte migration within the midbrain starting from 64hpf. In the hindbrain, *col1a2*+ve pericytes at 3dpf are always located on the MceV, extending their process into the hindbrain or adjacent to the large vessels around the meninges, just as in the midbrain (Fig. S5E-G).

**Fig. 5.**
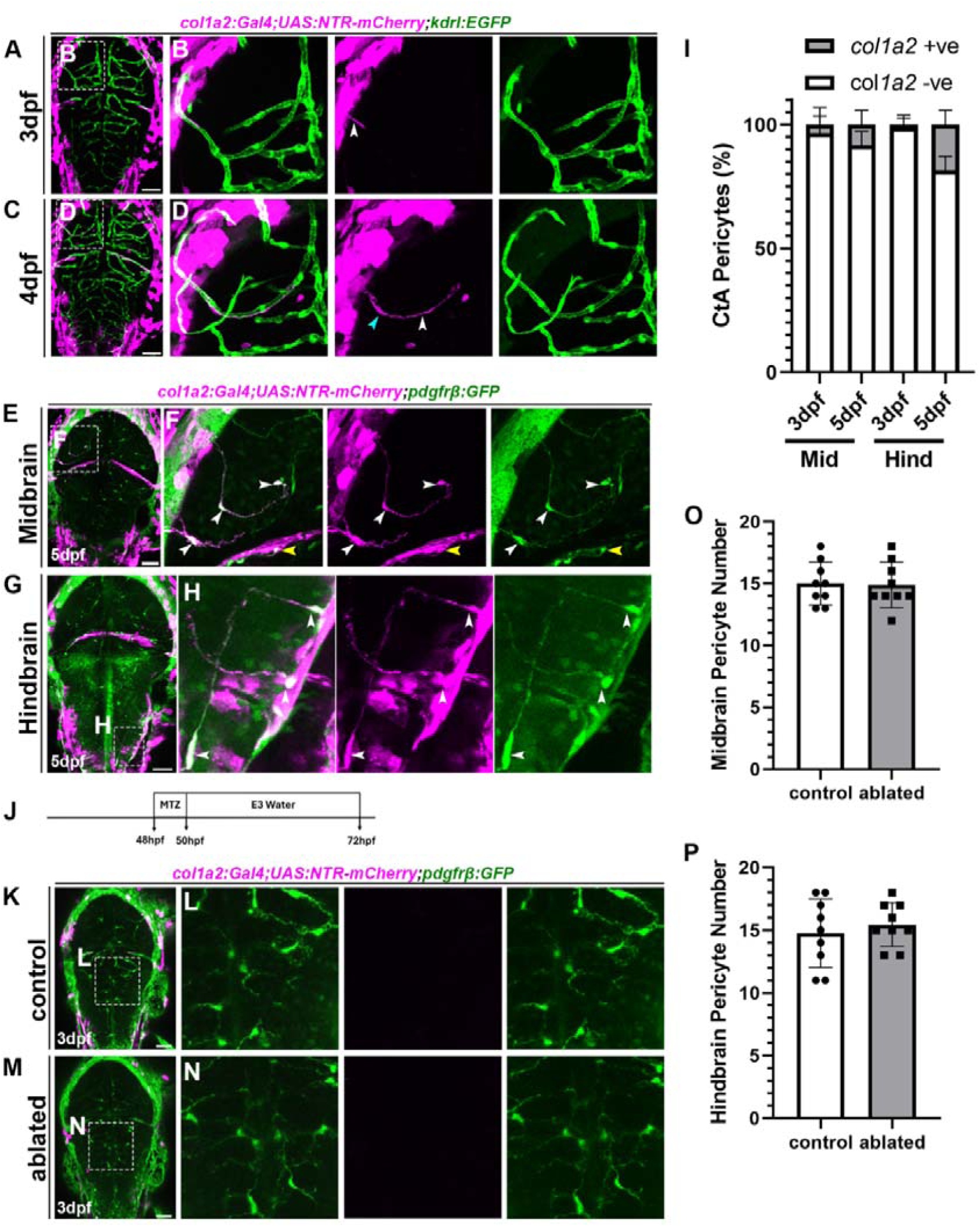
*col1a2*-expressing cells give rise to late embryonic brain pericytes. (A-D) Dorsal view of zebrafish larva at (A-B) 3dpf, (C-D) 4dpf on *col1a2* (magenta) and endothelial (*kdrl*; green) reporter lines. The insets show the pericyte process structure (white arrow) and soma (blue arrow). (E-H) Confocal image of 5dpf zebrafish larva *col1a2* and *pdgfr*β transgenic lines. The inset shows pericytes that co-express *col1a2* on the brain CtAs (white arrows) and Mcev (yellow arrow). (I) Quantification of CtA pericytes that co-express *col1a2* at 3 and 5dpf (n: 3dpf = 9, 5dpf =12). (J) Schematic of metronidazole (MTZ) drug treatment. (K-N) Dorsal view of a (K-L) control and (M-N) MTZ ablated 3dpf larva on *col1a2* and *pdgfr*β transgenic lines. The hindbrain inset image shows no co-expression of *col1a2* with *pdgfr*β, and no difference in pericyte number between the control and ablated groups. (O-P) Quantification of (O) midbrain and (P) hindbrain pericyte number between control and ablated group at 3dpf (n: control & ablated=9). Statistical analyses used unpaired t-tests, with results presented as mean ± SD. Scale bars: 50µm.

The behavior of *col1a2*-expressing cells is similar at 5dpf, as we observe *col1a2*+ve pericytes in similar locations, proximal to the meninges of the mid- and hindbrain and around the MceV (Fig. 5E-H). The proportion of *col1a2*+ve *pdgfr*β+ve pericytes at 3 & 5 dpf increases from 3.31% to 8.5% of midbrain and 1.31% to 18.5% of hindbrain pericytes (Fig. 5I). This suggests that *col1a2*+ve pericytes are a quantitatively significant source of pericytes migrating into the brain in late embryonic development. To demonstrate the meningeal origin of this cell, we used photoconversion of a *col1a2^kaede^*reporter. Upon exposure to UV light, *col1a2^kaede^* undergoes photoconversion, switching from green to red fluorescence. This will allow us to trace the migration of converted cells. We converted half of the brain at 55hpf (Fig. S6A-B, B’-B’’’). While the *col1a2* line is mosaic, most meningeal *col1a2-*expressing cells were converted (Fig. S6C-D, D’-D’’’). By 4dpf, we observe that *col1a2* meningeal cells from the midbrain region have entered the brain and assume a typical pericyte morphology (Fig. S6E-F, F’-F’’’). This suggests that *col1a2* lineage precursors migrate from the meningeal layer into the brain and differentiate into pericytes.

Since there are spatio-temporal differences between the behavior of *col1a2-*expressing and *col5a1-*expressing cells on the vessels, we wanted to test the effect of ablating early *col1a2-*expressing cells via metronidazole treatment from 48-50hpf (Fig. 5J). At 3dpf, there is no expression of *col1a2* in *pdgfr*β-positive hindbrain pericytes, located along the brain midline (Fig. 5K-N). Ablating *col1a2*-expressing cells has no effect on the number of pericytes, in contrast to the ablation of *col5a1*-expressing cells, where pericyte number was decreased (Fig. 5O-P vs. Fig. 4L-M). This suggests that early-migrating pericytes do not express *col1a2*.

It is important to distinguish whether the *col1a2* cell lineage (i.e., the transcriptional reporter) rather than the *col1a2* gene is important in late pericytes. We find no significant differences in pericyte number, vessel network length, pericyte coverage, or density between homozygous *col1a2^ca108^*mutants and their siblings (Fig. S7A-H). This suggests that while the lineage of cells expressing *col1a2* are pericyte precursors, the *col1a2* gene itself is dispensable for pericyte differentiation.

### *notch3* mutant pericytes have elevated *col1a2* mRNA expression

We next examined differences in *col5a1* and *col1a2* mRNA expression during normal development using hybridization chain reaction (HCR) RNA-FISH to label *col5a1*, *col1a2*, and *kdrl* (Fig. 6A-C). At 55 hpf, we analyzed the mid- and hindbrain CtAs and observed that only *col5a1* was expressed on the endothelium at this time, while *col1a2* was lacking in this region (Fig. 6B’-B’’’’, C’–C’’’’). At 5 dpf, when pericyte coverage is dense (Fig. 6D), we observe high expression of *col5a1* on the CtAs, while *col1a2* was nearly absent on these vessels (Fig. 6E’–E’’’’). *col1a2* expression is restricted to perivascular expression near the MceV, in addition to *col5a1* expression on the same vessel. This is consistent with our transgenic data, which showed that *col1a2* cells migrate and populate the MceV during development. However, unlike our transgenic reporter lines, where mCherry has strong persistence, and we can label *col1a2*-derived cells, our mRNA data show that this gene is turned off in the CtAs at both 55 hpf and 5dpf, and that only *col5a1* is still expressed at the start of pericyte migration and when pericytes are densely populated. Nonetheless, both *col5a1* and *col1a2* are expressed adjacent to the endothelium in the MceV. Also, in some CtAs, *col5a1* is co-expressed with *pdgfr*β, whereas in others, *pdgfr*β cells lack *col5a1* (Fig. 6G’–G’’’’). In summary, our RNA FISH data show that in normal development, *col5a1* mRNA expression is localized to the CtAs at 5dpf, while *col1a2* is not.

**Fig. 6.**
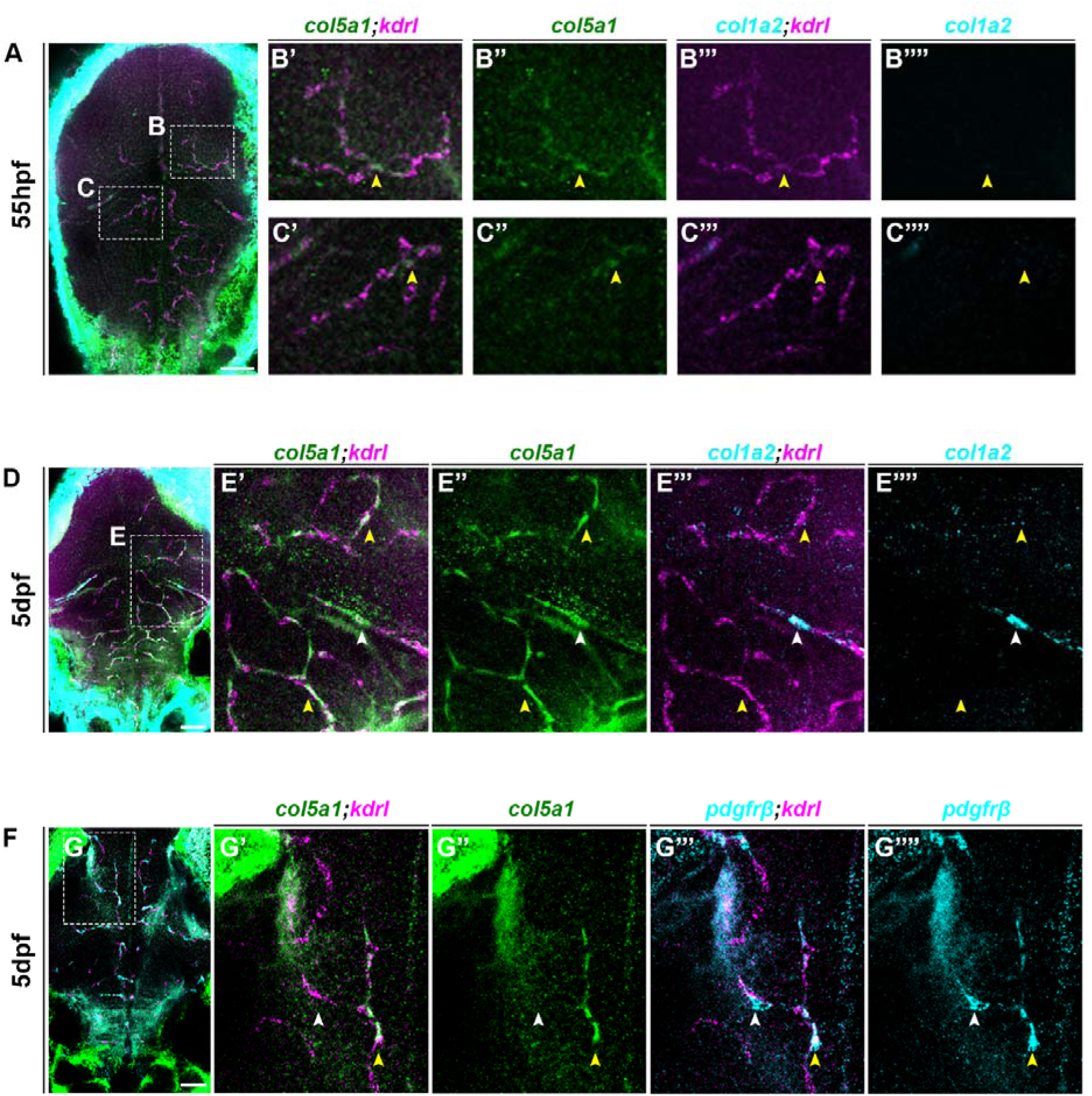
col5a1 mRNA is highly expressed on CtA vessels during embryonic development. (A-C) HCR mRNA in situ of *col5a1*, *col1a2*, and *kdrl* probes on a 55hpf larva. (B’-B’’’’) inset from the midbrain with yellow arrows indicating perivascular expression of *col5a1* and not *col1a2* on the vessels (kdrl). (C’-C’’’’) inset from the hindbrain with yellow arrows indicating CtAs where *col5a1* expression is evident (D-E) mRNA in situ of a 5dpf larva marked with *col5a1*, *col1a2*, and *kdrl* probes. (E’-E’’’’) Inset with yellow arrows indicates the CtAs are marked by *col5a1* expression but lack *col1a2* expression. White arrows indicate the MceV has both *col5a1* and *col1a2* gene expression. (F-G) mRNA in situ of a 5dpf larva marked with *col5a1*, *pdgfr*β, and *kdrl* probes. (G-G’’’’’) Inset with yellow arrow shows double positive staining for *col5a1* and *pdgfr*β, and the white arrow shows a *pdgfr*β positive cell that lacks *col5a1*. Scale bars: 50µm.

Thus, we investigated these gene expressions in the *notch3* small vessel disease model. In humans, *NOTCH3* homozygous loss-of-function mutations cause infantile and early-onset childhood stroke with cerebral white matter hyperintensities as seen on brain MRI images (Greisenegger et al., 2021; Pippucci et al., 2015; Stellingwerff, 2022). In zebrafish, loss of *notch3* causes intraventricular hemorrhage, a leaky blood-brain barrier, and reduced pericyte number at 3dpf (Wang et al., 2014). However, much remains unknown about pericyte characteristics in this *notch3* model. We observed reduced pericyte number and shorter vessel network lengths at both 3 and 5 dpf in homozygous *notch3^fh322^* mutants, consistent with previous observations (Fig. S8A-H). Interestingly, not only was the number of pericytes reduced in homozygous *notch3^fh322^* mutants, but we also observed that these pericytes have abnormal morphology with shorter process length (Fig. 7A-D, D’–D’’) in homozygous mutants, with an average length of 54.17µm compared with siblings at 86.52µm (Fig. 7E).

**Fig. 7.**
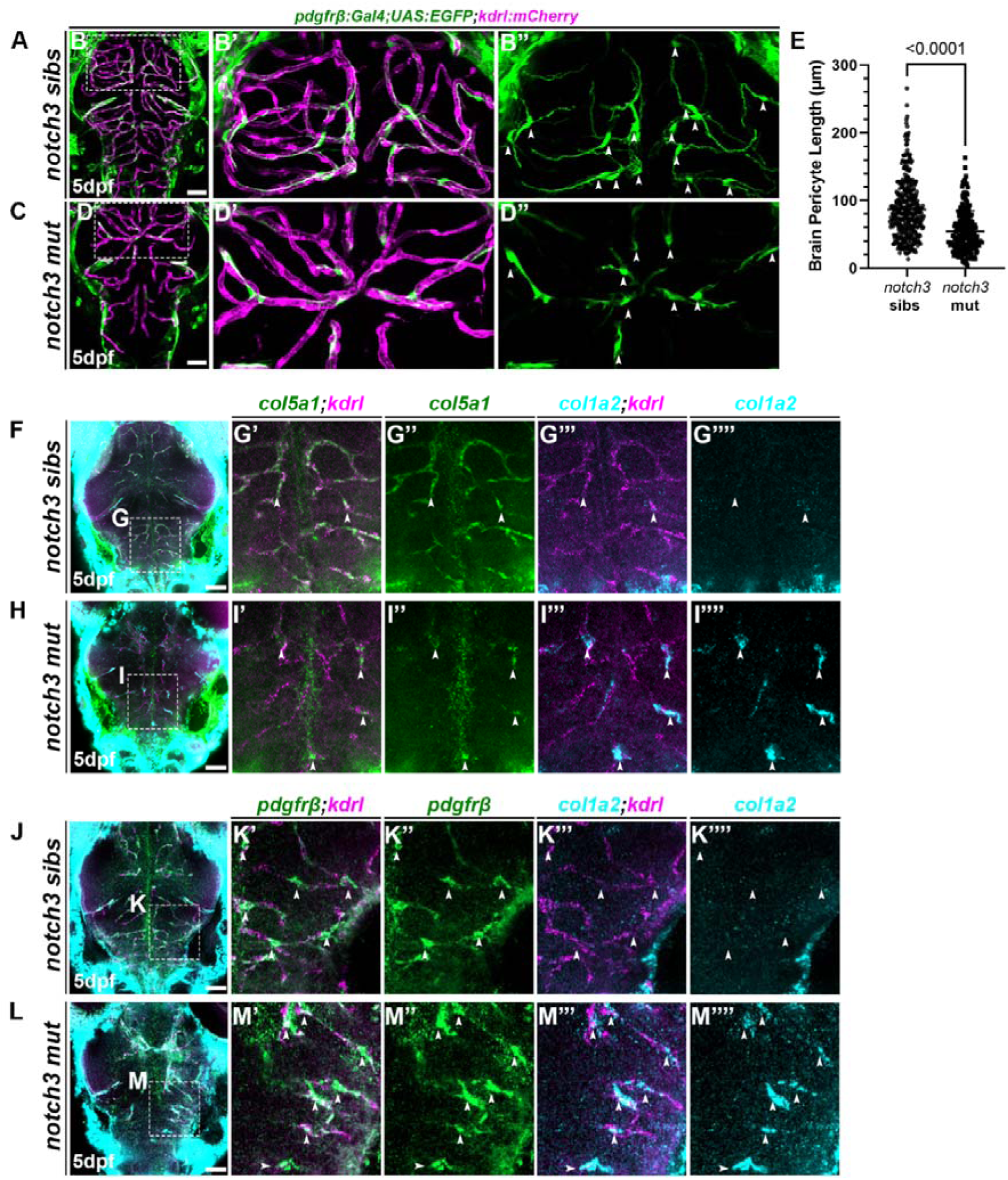
*notch3^fh322^*mutants pericytes have elevated *col1a2* expression. (A-D) Dorsal view of a (A-B*) notch3^fh322^* siblings (C-D) homozygous mutant at 5dpf on pericyte (*pdgfr*β; green) and endothelial (*kdrl*; magenta) transgenic lines. (B’-B’’) White arrows point to *notch3^fh322^*sibling pericytes in the image. (C’-C’’) White arrows points to homozygous mutant pericyte with shorter length or with unusual morphology (E) Quantification of brain pericyte length at 5dpf shows homozygous *notch3^fh322^* mutants pericytes are shorter (n: sibs= 7, mut= 10) (F-I) Dorsal view of an HCR mRNA in situ of *col5a1*, *col1a2*, and *kdrl* probes on a 5dpf (F-G*) notch3^fh322^* siblings and (H-I) homozygous mutant. (G’-G’’’’) Inset of *notch3^fh322^* siblings with white arrows indicating strong *col5a1* expression and no *col1a2*. (I’-I’’’’) Inset of homozygous *notch3^fh322^* mutants with white arrows indicating reduced *col5a1* and elevated *col1a2* expression on the vessels. (J-M) HCR mRNA in situ of *pdgfr*β, *col1a2*, and *kdrl* probes on a 5dpf (J-K*) notch3^fh322^* siblings (L-M) homozygous mutant. (K’-K’’’’) Inset of *notch3^fh322^* siblings with white arrows indicating CtA pericytes do not express *col1a2* (M’-M’’’’) Inset showing homozygous *notch3^fh322^* mutant pericytes express *col1a2*. Scale bars: 50µm.

Furthermore, as expected, *notch3^fh322^* siblings have strong *col5a1* expression on the CtAs at 5 dpf but lack *col1a2* (Fig. 7G’–G’’’’). On the other hand, homozygous *notch3^fh322^* mutants have reduced *col5a1* on the CtAs, but upregulated *col1a2* expression (Fig. 7I’–I’’’’). Thus, we investigated whether the elevated *col1a2* expression was in pericytes. We used HCR probes *pdgfrb* to label pericytes, along with *col1a2* and *kdrl* for gene expression analysis (Fig. 7J-M). We noticed that while the pericytes of *notch3^fh322^* siblings lacked *col1a2*, their homozygous mutants pericytes showed strong expression of *col1a2* (Fig. 7K’–K’’’’, M’–M’’’’). This *col1a2* staining showed pericytes with abnormal morphology and short processes, similar to the transgenic reporter. Upregulation of *col1a2* expression in the homozygous *notch3^fh322^* mutant suggests that pericytes may directly influence ECM remodeling, potentially inducing fibrosis in this SVD model. High collagen I is a characteristic of fibrosis and can lead to vessel stiffness and increased wall thickness (Fleenor et al., 2010).

## DISCUSSION

Pericytes have heterogeneous origins and phenotypes (Grant et al., 2019). How, where, and when pericytes from different origins enter the brain and contribute to subsequent heterogeneity have not been systematically examined. Our findings deepen understanding of the temporal and lineage dynamics of early brain pericyte development (Fig 8).

**Fig. 8.**
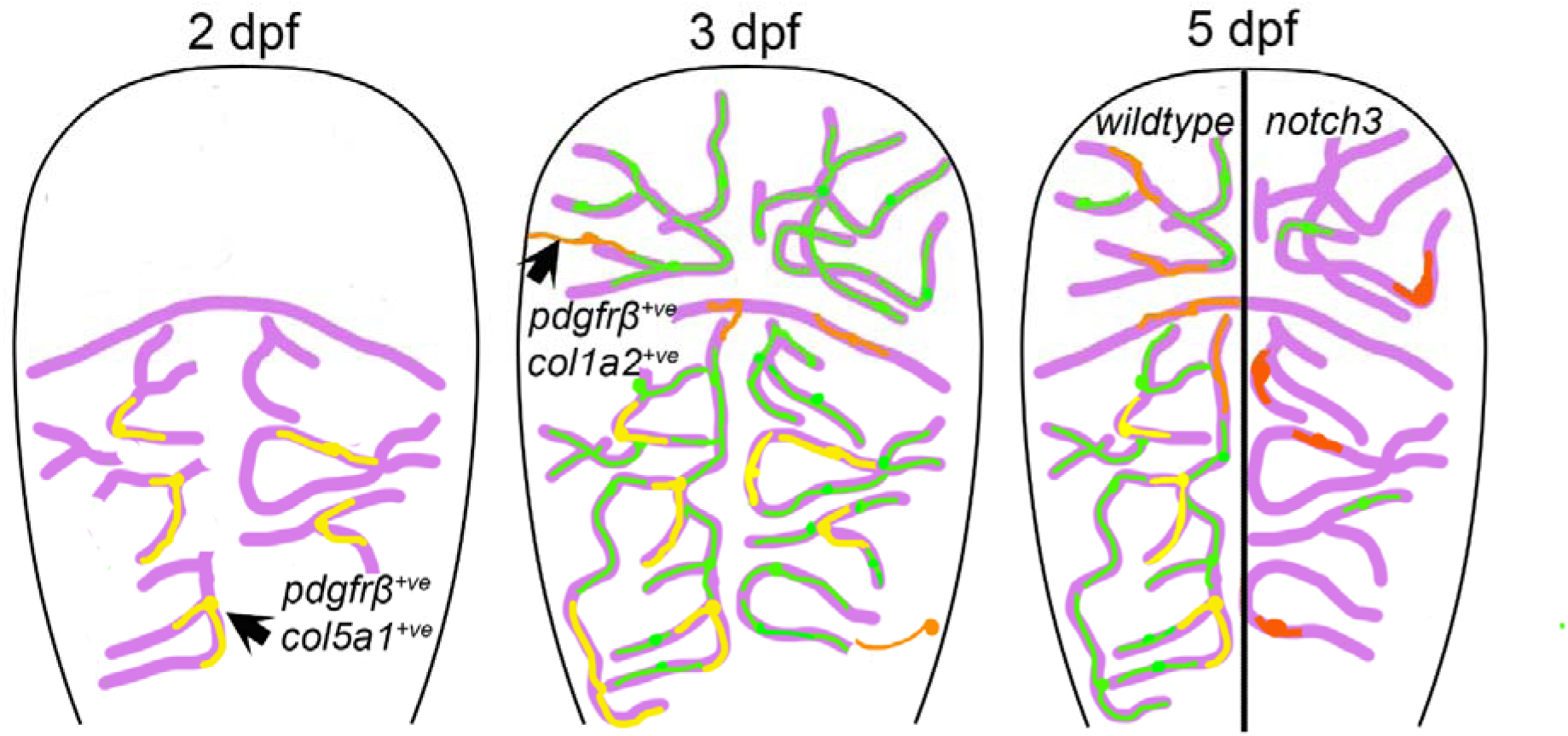
Model of progressive coverage of zebrafish brain vessels with pericytes. In early development, pericyte precursors expressing *pdgfr*β and *col5a1*(yellow) enter the brain from ventral sources and migrate along vessels (purple). At 3 dpf, there is some proliferation to expand the population, and most cells express both *pdgfr*β and *col5a1 (yellow),* although a small number of *pdgr*β*-*only expressing cells are also present (green). Cells expressing *col1a2* extend processes towards brain vessels (orange) at this stage. By 5 dpf, cells expressing a mixture of markers are established on the brain vasculature in wildtype. In *notch3* mutants, there are fewer *pdgfr*β-expressing cells, reduced *col5a1,* and increased and abnormal *col1a2* expression (red).

Zebrafish brain pericytes are mostly associated with capillaries, such as the small-caliber Central Arteries (CtAs) formed via angiogenesis, which have a comparable diameter to mammalian capillaries (Bahrami and Childs, 2020; Whitesell et al., 2019). Our study demonstrates the rapid pace at which pericytes must develop to keep up with endothelial expansion, as we observed an approximately 200% increase in CtA pericyte number and a 30-70% increase in pericyte coverage from 3 to 5 dpf. We showed that pericyte recruitment is sensitive to angiogenesis, as reducing it via the general Vegfr or brain-specific Gpr124 signaling reduces pericyte number and coverage. This reduced pericyte coverage in the angiogenesis-inhibited group suggests that pericyte-endothelial interactions with a healthy endothelium are needed to support pericyte process growth during development. Extracellular matrix proteins, such as fibronectin, which are deposited abundantly during angiogenesis, have been implicated as the preferred matrix for pericyte process migration (Dessalles et al., 2021; Hielscher et al., 2016). Fibronectin binds to the integrin α5β1 heterodimer, and specific deletion of *Itgb1* (integrin β1) in pericytes results in shorter cytoplasmic processes (Abraham et al., 2008). While our study highlights the importance of a healthy endothelium for pericyte growth and process extension, more research is needed to fully understand the role of these ECM proteins in pericyte expansion and cytoplasmic process extension, especially during angiogenesis.

Brain pericytes first originate from cell differentiation around 2dpf, a switch from *pdgfr*β*^low^* to *pdgfr*β*^high^*, migrating cells from the large vessels to the CtAs, then proliferate thereafter (Ando et al., 2016; Ando et al., 2019). Proliferation has been suggested as a major mechanism by which pericytes expand their population. In cell culture studies, pericytes co-cultured with endothelial cells divide more around sprout elongation and have enhanced proliferation, with about 88% positive for the S phase EdU proliferative marker (Chiaverina et al., 2019; Payne et al., 2021; Tarallo et al., 2012). In vivo, additional mechanisms, like increased blood flow, can result in an increase in embryonic brain pericyte number and vessel coverage via activation of Piezo-Notch signaling (Zi et al., 2024). Using the fluorescent ubiquitination-based cell cycle indicator (FUCCI) system, this study labelled about 20% of pericytes in S/G2/M phase at 3.5dpf (Zi et al., 2024). Our findings, obtained using both time-lapse imaging and proliferation markers that label the S/M Phase, suggest that in vivo, embryonic pericytes have low proliferation rates, with about 8.6% in the S Phase. The low proliferation rate suggests that continuous cell differentiation is needed to support brain pericyte number expansion from pericyte precursors that are present around larger-caliber vessels (like the basilar artery) before migrating and differentiating to the mid- and hindbrain CtAs (Ando et al., 2021), although this likely still doesn’t account for the increase in pericytes in late embryogenesis in zebrafish.

Previous work from our lab has shown that the transcription factor *nkx3.1* regulates brain pericyte differentiation, and it is enriched in fibroblasts expressing *col1a2* and *col5a1* (Ahuja et al., 2024; Rajan et al., 2023). Both *col5a1* and *col1a2* are expressed in trunk pericyte progenitors. Therefore, we did not expect cells from the transgenic lines reporting *col5a1* and *col1a2* expression to behave differently, but we found both spatial and temporal differences. *col5a1-expressing* precursors are present in the basilar artery and migrate to the CtAs, to give rise to the first set of pericytes, similar to what is seen for *pdgfr*β-expressing pericytes (Ando et al., 2016). These cells appear around the midline of the hindbrain by 48hpf. To highlight the importance of these first wave pericytes, ablation of *col5a1+ve* cells from 48–50hpf results in a regional decrease of brain pericytes, particularly in the hindbrain CtAs along the midline. However, we were surprised by the contrasting behavior of *col1a2*+ve pericytes, which are first seen at 64hpf in the midbrain. *col1a2*+ve pericytes in the midbrain arise from the larger vessels around the meninges, while those in the hindbrain migrate from the midbrain cerebral vein (MceV) and the primordial hindbrain channel (PHBC). Our findings show that distinct fibroblast precursor lineages are present on large brain vessels that migrate and differentiate into CtA pericytes at different times during development.

In mice, *Col1a1*/*Col1a2-*expressing fibroblasts develop embryonically in the meninges but do not associate with the brain vessels until postnatal development (DeSisto et al., 2020; Kelly et al., 2016). They migrate and proliferate around the ascending arterioles & venules after vascular smooth muscle cells and astrocytes’ end feet are established (Jones et al., 2023). These perivascular fibroblasts express the pericyte marker *Pdgfr*β and develop concurrently with the perivascular macrophages along the cortical vessels (Jones et al., 2023; Jones et al., 2026). In a possible parallel, we observed *col1a2*+ve *pdgfr*β*-*expressing cells with pericyte morphology on the smaller vessels emerging in the midbrain at 64hpf. Their time of emergence matches when the early embryonic zebrafish brain macrophages become mature, and transition to downregulate *L-plastin* and upregulate *apolipoprotein E* (*apo E*) between 60-72hpf (Herbomel et al., 2001). Zebrafish vascular smooth muscle cell differentiation in the head starts around 72hpf with robust expression between 96-120hpf (Whitesell et al., 2014; Whitesell et al., 2019). Similarly, we observe an increase in the populations of pericytes co-expressing *col1a2* from 3.31% of mid and 1.31% of hindbrain pericytes at 3dpf to about 8.5% of mid and 18.5% of hindbrain pericytes at 5dpf. The developmental timing of pericyte emergence is consistent with that of other cells in the neurovascular unit.

Furthermore, we find that *col5a1* is preferentially expressed on the small-caliber CtAs vessels, while *col1a2* is preferentially expressed on the MceV. During vascular injury, ECM remodeling switches from elastin to a collagen-rich deposition (Lin and Davis, 2023). We tested *notch3^fh322^* mutants that hemorrhage and are pericyte-deficient to show that *col5a1* (type V collagen) is downregulated in the CtAs, with a pronounced upregulation of *col1a2* (type I collagen). Under homeostatic conditions, brain fibroblast cells are important for maintaining vascular structural integrity by depositing collagen; however, overexpression of collagen I in pathological conditions can lead to fibrosis (Duan and Yu, 2024). In the *notch3^fh322^* mutants, *col1a2* is upregulated in pericytes with abnormal morphology. Although changes in pericytes to a more fibrotic phenotype in vivo are not well characterized, vascular smooth muscle cells undergo a phenotypic switch to myofibroblasts, which thicken the vessel wall during injury or in disease (Lu et al., 2020). Following arterial injury, *Col1a2* is activated via SRF/SMAD3-TGFβ signaling in vascular smooth muscle cells, which causes vascular fibrosis (Shen et al., 2024). While we have shown a similar phenotypic switch with elevated *col1a2* expression in *notch3^fh322^* mutant pericytes, further work is needed to determine whether TGFβ signaling plays a role in this.

## MATERIALS AND METHODS

### Zebrafish Husbandry and Transgenic Lines

All experimental protocols were approved by the University of Calgary’s Animal Care Committee (AC25-0151). Zebrafish embryos were maintained in an incubator at 28.5°C in E3 media. For experiments that tracked the same embryo across development, each zebrafish larva was transferred to a well of a 6-well plate after initial imaging. From 5dpf onward, the larvae were fed rotifers daily, with daily changes of E3 water (5ml), until the time of analysis, with a maximum duration of 10dpf. The zebrafish lines include: *TgBAC(pdgfrb:Gal4FF)^ca42^* (Whitesell et al., 2019),*Tg(kdrl:mCherry)^ci5^* (Proulx et al., 2010), *TgBAC(pdgfrb:GFP)^ca42^* (Whitesell et al., 2019), *TgBAC(col1a2:Gal4-VP16)^ca102^* (Ma et al., 2018), *Tg(kdrl:EGFP)^la116^* (Choi et al., 2007), *Tg(top:GFP)^w25^* (Dorsky et al., 2002), *Tg(UAS-NTR:mCherry)^c264^* (Davison et al., 2007), *Tg(5xUAS :EGFP)^nkuasgfp1a^* (Asakawa et al., 2008), *Tg(UAS:kaede)^s1999t^* (Davison et al., 2007), *col1a2^ca108^* (Rajan et al., 2020), *notch3^fh322^* (Alunni et al., 2013).

### Generation of TgKI(col5a1:2A-Gal4-VP16)^ca120^ line

The *TgKI(col5a1:2A-Gal4-VP16)^ca120^* line was generated following the GeneWeld protocol (Welker et al., 2021; Wierson et al., 2020). Briefly, synthesized *col5a1* crRNA (5’-GTTGGACCCTCAGGACCGGA-3’), universal crRNA (5’-GGGAGGCGTTCGGGCCACAG-3’), and tracrRNA were ordered from IDT. A donor plasmid containing the 2A-Gal4-VP16 cassette with 48 bp of homology arms corresponding to the *col5a1* sequence, flanked by a universal CRISPR crRNA target sequence, was generated. To generate knock-in lines, 1 nL of an injection mix containing 100 ng/µL Cas9 mRNA, 5 µM *col5a1* crRNA:tracrRNA duplex, 5 µM universal crRNA:tracrRNA duplex, and 10 ng/µL donor plasmid was co-injected into *UAS:NTR-mCherry* embryos at the one-cell stage. mCherry^+^ embryos were raised to adulthood. Stable knock-in lines were established by screening F1 embryos from injected founders for mCherry expression. In this manuscript, the *TgKI(col5a1:2A-Gal4-VP16)^ca120^; Tg(UAS-NTR:mCherry)^c264^* cross is referred to as col5a1*^NTR-mCherry^*.

### Embryo Microinjection, Drug Treatment, and Genotyping

Gpr124 morpholino (Table S1) was injected into one-cell stage zebrafish embryos. The list of drugs and their working concentration is available in Table S2. Drug treatments were done in a 24-well plate with 10 – 15 dechorionated embryos per well. DMH4 and SU5416 were dissolved in DMSO to prepare stock solutions at 10 mM and 1mM, respectively. Stock solutions were diluted to the desired working concentration in E3. For metronidazole, the stock solution 100mM and the working solution 5mM were prepared in E3. Sibling embryos in an equivalent dilution volume of DMSO or E3 were used as controls. *col1a2^ca108^* embryos were genotyped as previously described by (Rajan et al., 2020). *notch3^fh322^* embryos were genotyped using a customed designed TaqMan SNP genotyping assay (ID: ANXG746, ThermoFisher).

### Antibody Staining and EdU assay

For pHH3 antibody staining, embryos were fixed in 4% paraformaldehyde for 2hrs at RT or overnight at 4°C, then washed for 3×5 mins in PBS 0.1% tween (PBT), followed by washes of 5mins each in a series of Methanol/PBT dilution (25/75%, 50/50%, 75/25%, 100/0%) and stored in 100% methanol overnight. Embryos were rehydrated with a series of Methanol/PBT (75/25%, 50/50%, 25/75%, 0/100%) 5-min washes, followed by 2×5-min PBT and 1×5-min PBS with 2% Triton X-100. Embryos were permeabilized with PBS 2% Triton X-100 for 2.5hrs at RT, then blocked for 1 – 2 hrs at 37°C °C in blocking solution (10% non-specific sheep serum, 1% DMSO, PBS 0.8% Triton X-100). Primary antibodies dilutions are in Table S3, and were incubated for 2 nights at 37°C. Samples were then washed in PBT for 6 × 15 min, the secondary antibody dilution was added, and the tubes were incubated in the dark at RT for 2 nights. Finally, samples were washed in PBT for 6 × 10 min and mounted for confocal microscopy imaging.

The EdU cell proliferation assay (Invitrogen C10338) was performed according to the manufacturer’s instructions, with minor modifications. About 15 – 20, 50hpf embryos were incubated in 100µl of 1mM EdU on an ice slurry for 30mins and returned to E3 water to grow until 80hpf. Embryos were fixed in 4% PFA for 2hrs at RT, followed by 3 × 5-min PBT washes, and then stored in PBT at 4°C overnight. Embryos were permeabilized in PBS containing 2% Triton X-100 for 2hrs, incubated in 3% BSA/PBS containing 0.8% Triton X-100 for 1hr at RT, followed by 3×5mins PBT washes. The click-it reaction cocktail was then added to embryos in tubes covered with tin foil and incubated at RT for a minimum of 3 hrs, followed by PBT washes for 3X5 mins and antibody blocking for 1hr at RT. Both primary and secondary antibodies were diluted appropriately and incubated overnight at 37°C, followed by final washes in PBT and mounting for image analysis.

### Hybridization Chain Reaction

Custom-made Hybridization Chain Reaction (HCR) probes were obtained from Molecular Instruments, Inc. (Los Angeles, California). Zebrafish whole-mount HCR was modified from the manufacturer’s instructions. Embryos were fixed in 4% PFA, 2hrs at RT for 55hpf or 2 nights at 4°C for 5dpf larvae, then washed for 3 × 5 min in PBT, followed by 5 min each in a series of Methanol/PBT dilutions (25/75%, 50/50%, 75/25%), and 2hrs of incubation in 100% methanol before storing overnight at 4 °C. Embryos were rehydrated with a series of Methanol/PBT (75/25%, 50/50%, 25/75%, 0/100%) 5-min washes, followed by 2 x 5-min PBT, and proteinase K digestion for 30 mins (40 µg/ml for 55hpf and 120 µg/ml for 5dpf). The embryos were rinsed in PBT and fixed in 4% PFA for 15 min, followed by 3 × 5-min PBT washes. For probe hybridization, embryos were incubated for 2 nights at 37°C. During the amplification phase, embryos were incubated for 1-3 nights at room temperature, the longer the incubation, the better the signal output. Reagents for HCR Gold RNA-FISH protocols were used with both HCR Gold and HCR v3 probes. The list of HCR probes and dilutions is available in Table S4.

### Confocal Microscopy

Zebrafish embryos were embedded in 0.8% low melting agarose (LMA) and fluorescent images were acquired using Zeiss LSM700/900 or 880 inverted with AiryScan (objective lens: Plan-Apochromat 20x/0.8 M27) with a slice interval of 1 – 1.75µm. For photoconversion of Kaede, the area of interest for was marked using the region tool. A timed bleaching experiment of 40-80 cycles was performed with the 405 nm laser. After photoconversion, embryos were returned to E3 water and re-imaged at 4dpf.

### Pericyte Measurements Image Analysis

Brain pericyte number was determined by splitting the confocal images into two z-stack projections containing one-half of the brain image. Subsequently, the multi-point tool in Fiji (v1.54f) was used to count cells on the CtAs that matched pericyte morphology, specifically soma body and process length. For pericyte process length, a smaller sub-z-stack with elaborate pericyte coverage was made (one z-stack can have 1-8 pericytes). The free-hand tool was used to longitudinally measure the pericyte length from the beginning of the soma to the end of the process. Vessel network length was automatically generated with the Python script ‘Vessel Metrics.’ (McGarry et al., 2024) (Fig. S1A-B). The vessel segments were output in pixels and then converted to microns based on the original confocal (.czi) image size. All non-brain (mid + hind) vessel segments were manually removed before calculating the total vessel network length, which is the sum of each brain vessel segment.

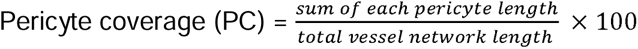

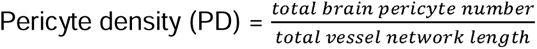

### Statistical Analysis

All statistical analyses were performed using GraphPad Prism v9.4.1 with all graphs plotted as mean ±SD and statistically significant defined as P value <0.05. A two-tailed unpaired t-test was used to analyze two-group comparison data and one-way ANOVA with Tukey’s test for multiple comparisons.

## ACKNOWLEDGEMENTS

The authors thank Pia Svendsen of the ACHRI imaging facility for microscopy training. We also thank C.U.A.’s supervisory committee members (Peng Huang, Maja Tarailo-Graovac), Sarah Childs, and members of Peng Huang’s lab for fruitful project discussions and feedback. Zebrafish schematics were created using https://www.biorender.com.

## Data and resource availability

All zebrafish lines are available at the Zebrafish Resource Center or available on request.

---

**Fig. S1.**
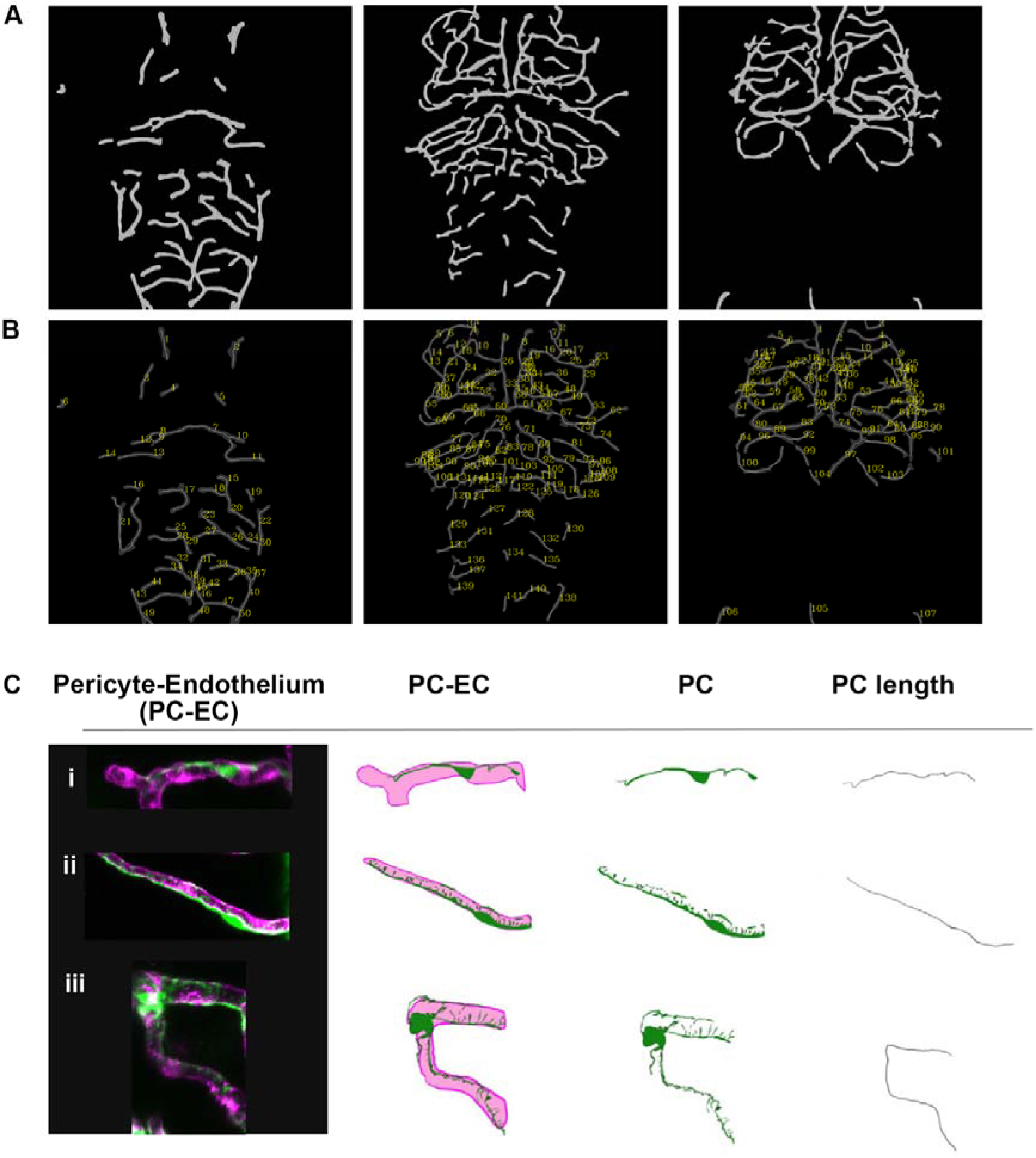
Measurement of vessel network and pericyte length. (A) Vessel Metrics (McGarry et al., 2024), segmentation of a 5dpf zebrafish larva in 3 sub-stacks of a Z-projection from the most dorsal to the ventral side of the brain. (B) Vessel Metrics numbered vessel segments corresponding to the segment vessel length in the .csv output file. (C) Three examples of pericytes on the brain endothelium showing how the longitudinal axis of the pericytes was measured for analysis.

**Fig. S2.**
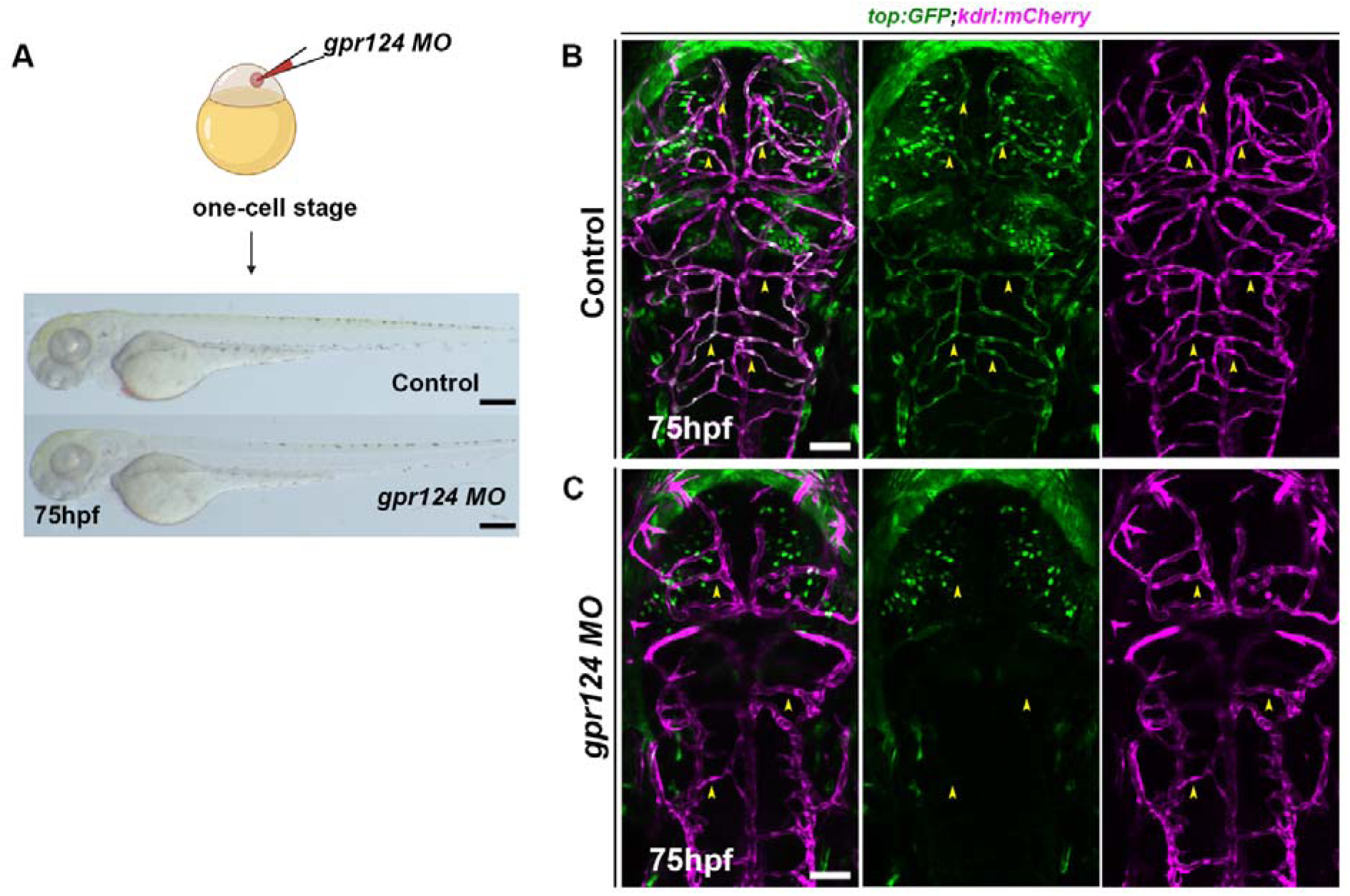
*gpr124* morpholino embryos lack Wnt activity in brain CtAs. (A) Schematic of *gpr124* morpholino injection at the one-cell stage and lateral image of 75hpf control and *gpr124* MO embryos indicating no morphological differences between groups. (B-C) Confocal image of 75hpf (B) Control and (C) *gpr124* MO larvae on Wnt reporter (green) and endothelial (magenta) transgenic lines. Yellow arrows show the Wnt activity on control CtAs and the lack of Wnt activity in CtAs of the *gpr124* MO group. Scale bars: 50µm.

**Fig. S3.**
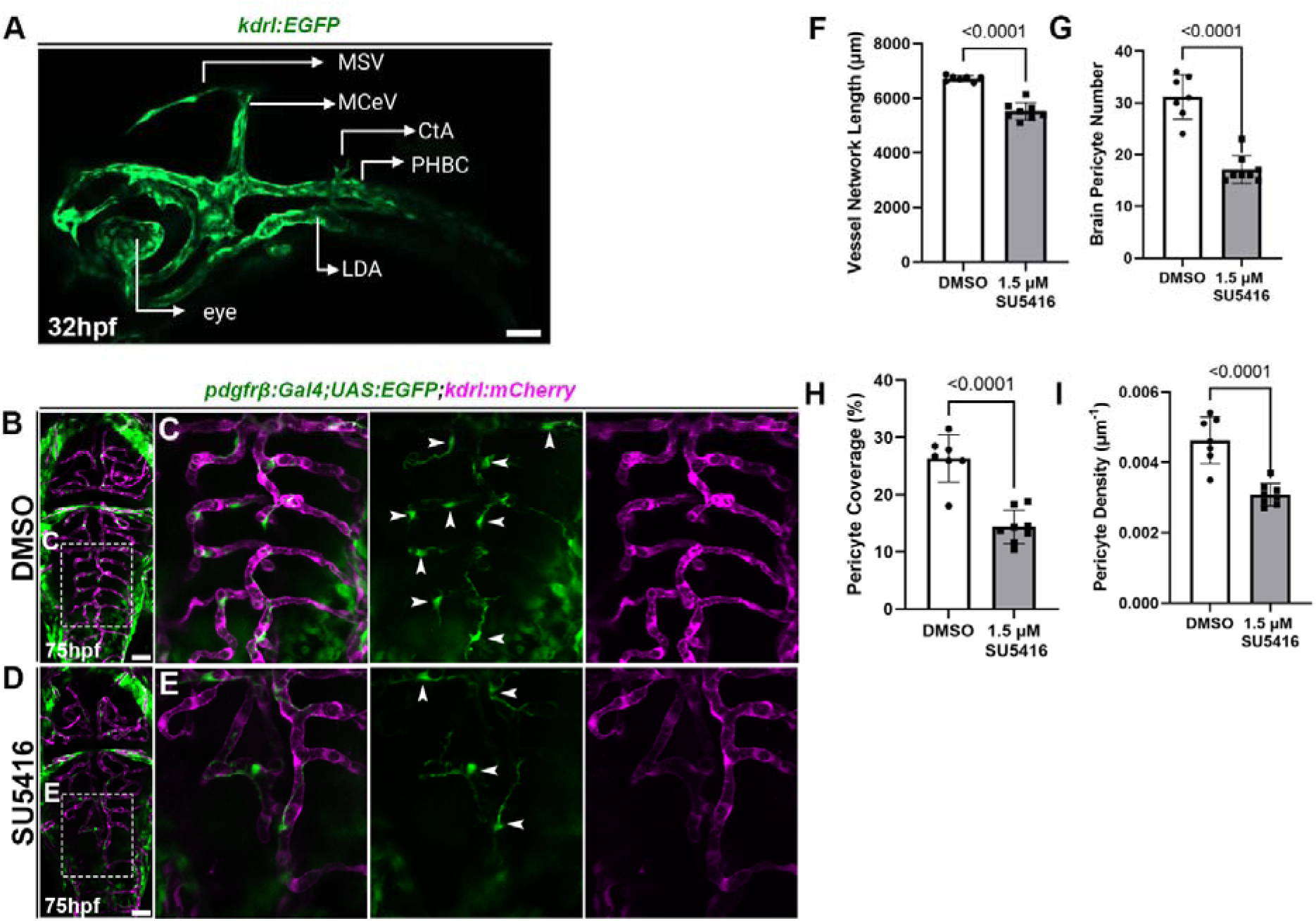
Reduced angiogenesis via Vegfr inhibition decreases pericyte numbers. (A) Lateral brain image of zebrafish embryo on endothelial line (green) at 32hpf shows vessel anatomy at the start of hindbrain CtA angiogenesis. (B-E) Dorsal view of 75hpf embryos on pericyte (green) and endothelial (purple) transgenic treated with (B-C) DMSO (D-E) SU5416 (Vegfr1/2 inhibitor). White arrows on hindbrain insets point to pericytes. (F-I) Quantification showing significantly reduced (F) vessel network length, (G) pericyte number, (H) pericyte coverage, and (I) pericyte density in DMH4-treated groups (n: DMSO=7, SU5416=8). LDA – lateral dorsal aorta, PHBC – primordial hindbrain channel, MceV – mid cerebral vein, MsV – mesencephalic vein. Statistical analyses are unpaired t-tests represented as mean ± sd. Scale bars: 50µm.

**Movie 1 Time-lapse movie of hindbrain pericyte proliferation and migration.** Two pericytes divide (white and yellow arrows) while new pericytes migrate onto the vessels (blue arrows).

**Fig. S4.**
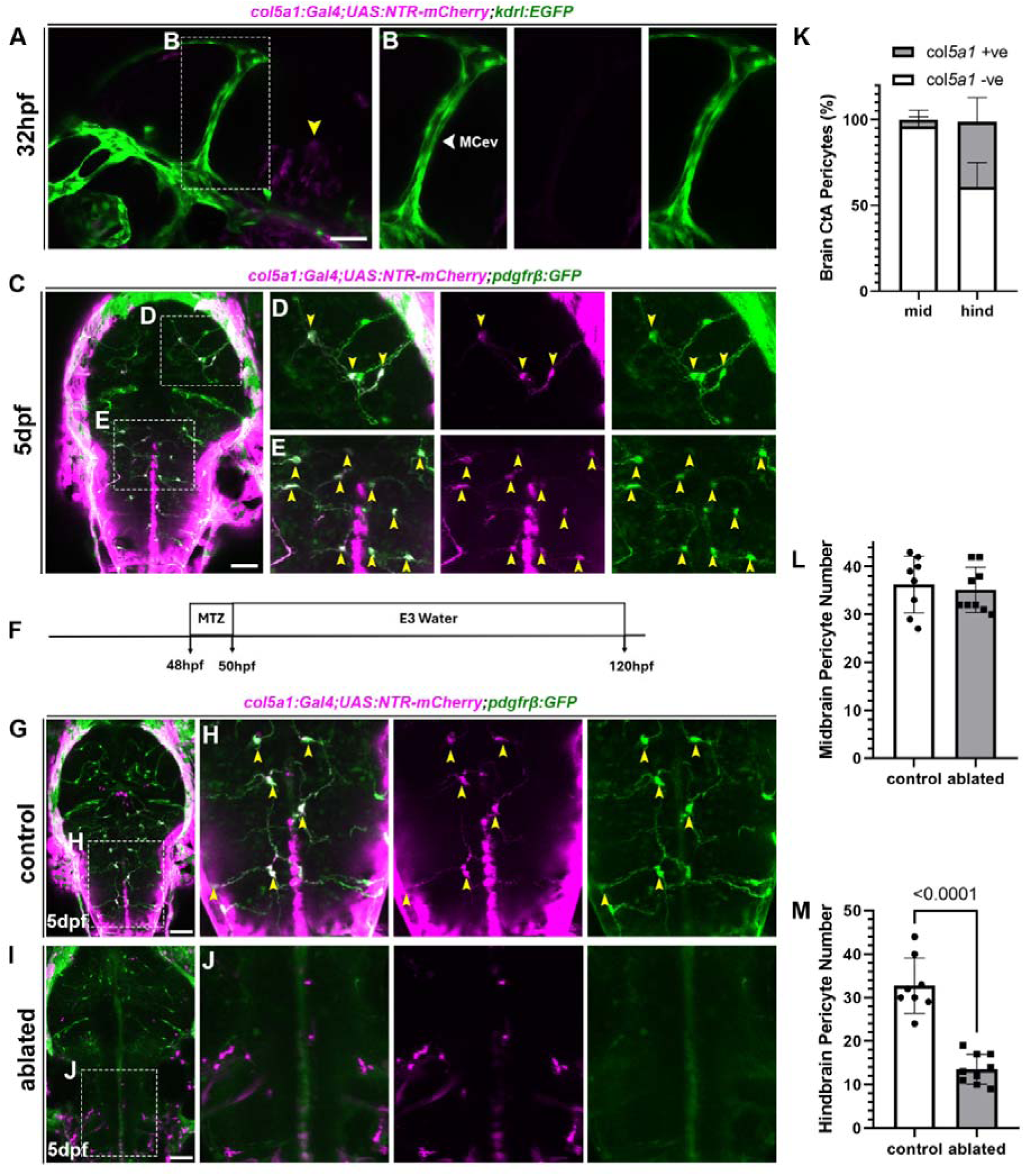
*col5a1*+ve pericytes persist through late embryogenesis. (A-B) Lateral view of a 32hpf zebrafish embryo on *col5a1* (magenta) and endothelial (green) reporter line. The yellow arrow points to *col5a1*+ve cells in the brain, while the white arrow shows *col5a1+ve* cells are absent from the MceV at 32hpf. (C-E) Zebrafish larva at 5dpf on *col5a1* and *pdgfr*β reporters. The yellow arrows in the inset indicate co-expression. (F) Schematic of metronidazole (MTZ) treatment and imaging timepoint. (G-J) Dorsal view of a (G-H) control and (I-J) MTZ-ablated 5dpf larva. Yellow arrows in the inset show *col5a1*+ve pericytes. (K) Graph showing the proportion of CtA pericytes that are positive or negative for *col5a1* expression at 5dpf (n=10). (L-M) Quantification of (L) midbrain and (M) hindbrain pericytes between control and MTZ-ablated groups (n: control =8, ablated =9). Statistical analyses are unpaired t-tests represented as mean ± sd. Scale bars: 50µm.

**Movie 2 *col5a1* cell migration from 48hpf** *col5a1* pericyte-like cells migrate and divide (yellow arrows) along the hindbrain CtAs

**Movie 3 Early *col1a2* cell migration** *col1a2* cells (magenta) migrate into the brain and associate with the mid cerebral vein (yellow arrow).

**Movie 4 Time of *col1a2*+ve pericytes appearance.** *col1a2*-ve pericytes migrate into the brain first (yellow arrows), before *col1a2*+ve pericyte emergence in the midbrain (white arrow).

**Fig. S5.**
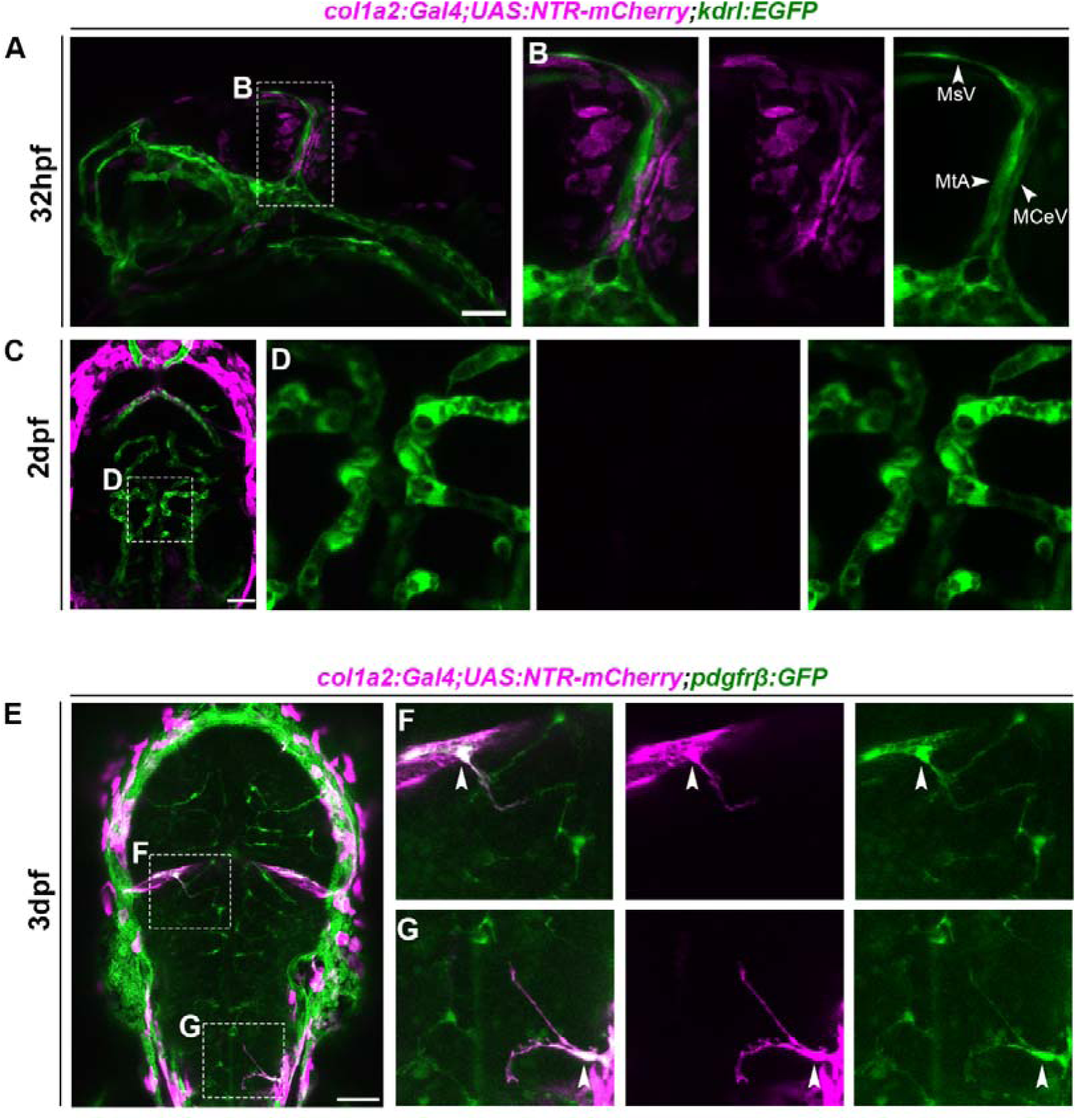
*col1a2* expressing cells first associate with large vessels before differentiating to pericytes. (A-B) Lateral view of zebrafish embryo at 32hpf on *col1a2* (magenta) and endothelial (*kdrl*; green) reporter line with *col1a2* fibroblast cells surrounding the MtA, MCeV, and MsV. (C-D) 2dpf zebrafish larvae with inset showing no *col1a2* +ve cells on the hindbrain CtAs. (E-G) Dorsal view of a 3dpf zebrafish larva on *col1a2* and *pdgfr*β transgenic lines. The inset image shows a pericyte migrating from the MCeV or the side of the hindbrain co-expressing the *col1a2* marker (white arrows). MtA – Metencephalic artery, MCeV- mid-cerebral vein, MsV- Mesencephalic vein. Scale bars: 50µm

**Fig. S6.**
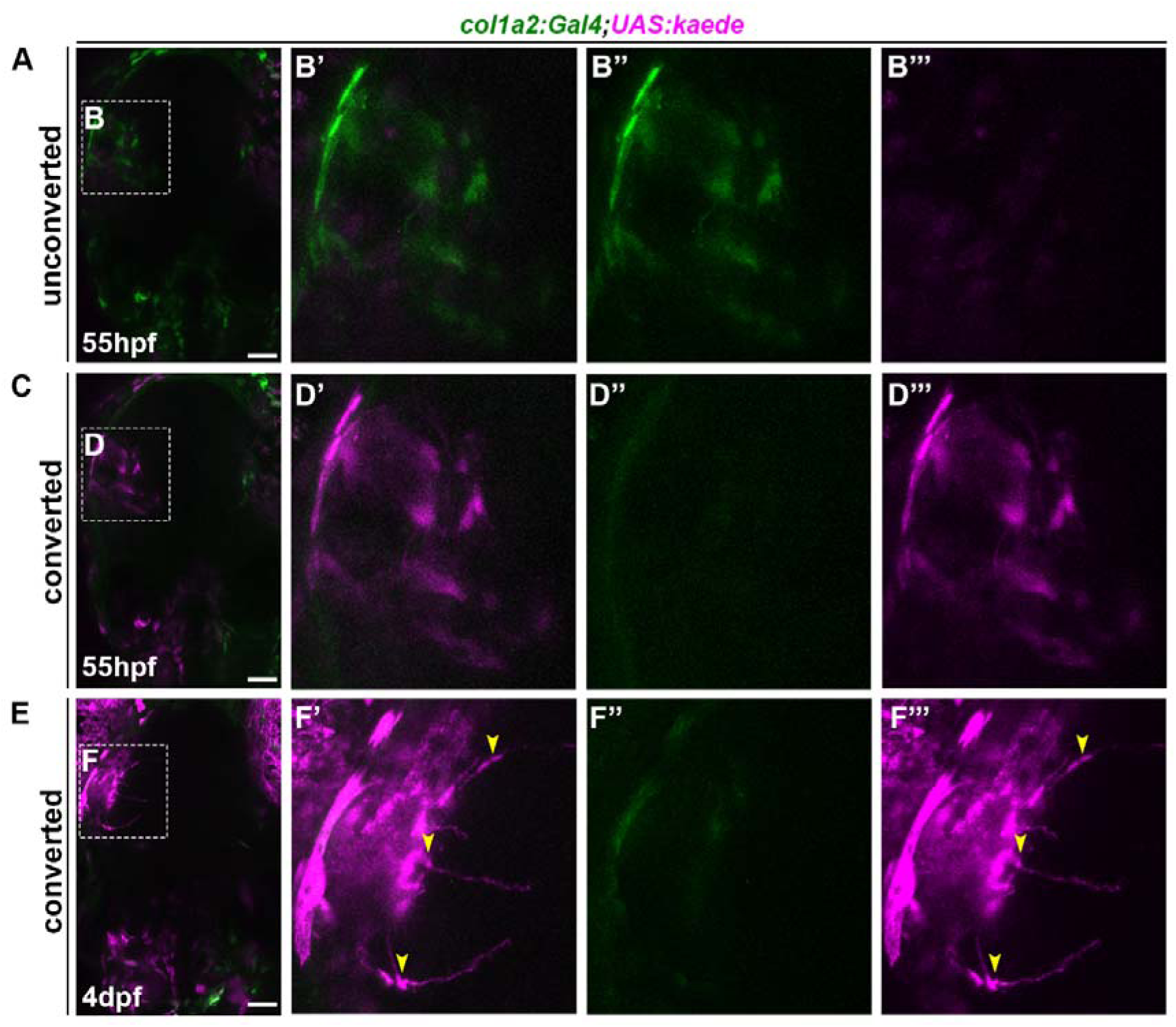
*col1a2* lineage tracing using Kaede photoconversion confirms *col1a2*-expressing cells become pericytes in the brain. (A-F) Confocal image of an embryo at 55hpf (A-B), before Kaede photoconversion (C-D), after Kaede photoconversion, and at 4pf (E-F) after Kaede photoconversion. (B’-B’’’) Area of interest of the midbrain shows *col1a2-*Kaede-expressing cells in green before photoconversion at 55hpf. (D’-D’’’) The area of interest after photoconversion shows that the green cells have converted to magenta at 55 hpf. (F’-F’’’) The inset shows an area of interest in the midbrain, with yellow arrows pointing to pericyte-like cells at 4dpf (n=3). Scale bars: 50µm.

**Fig. S7.**
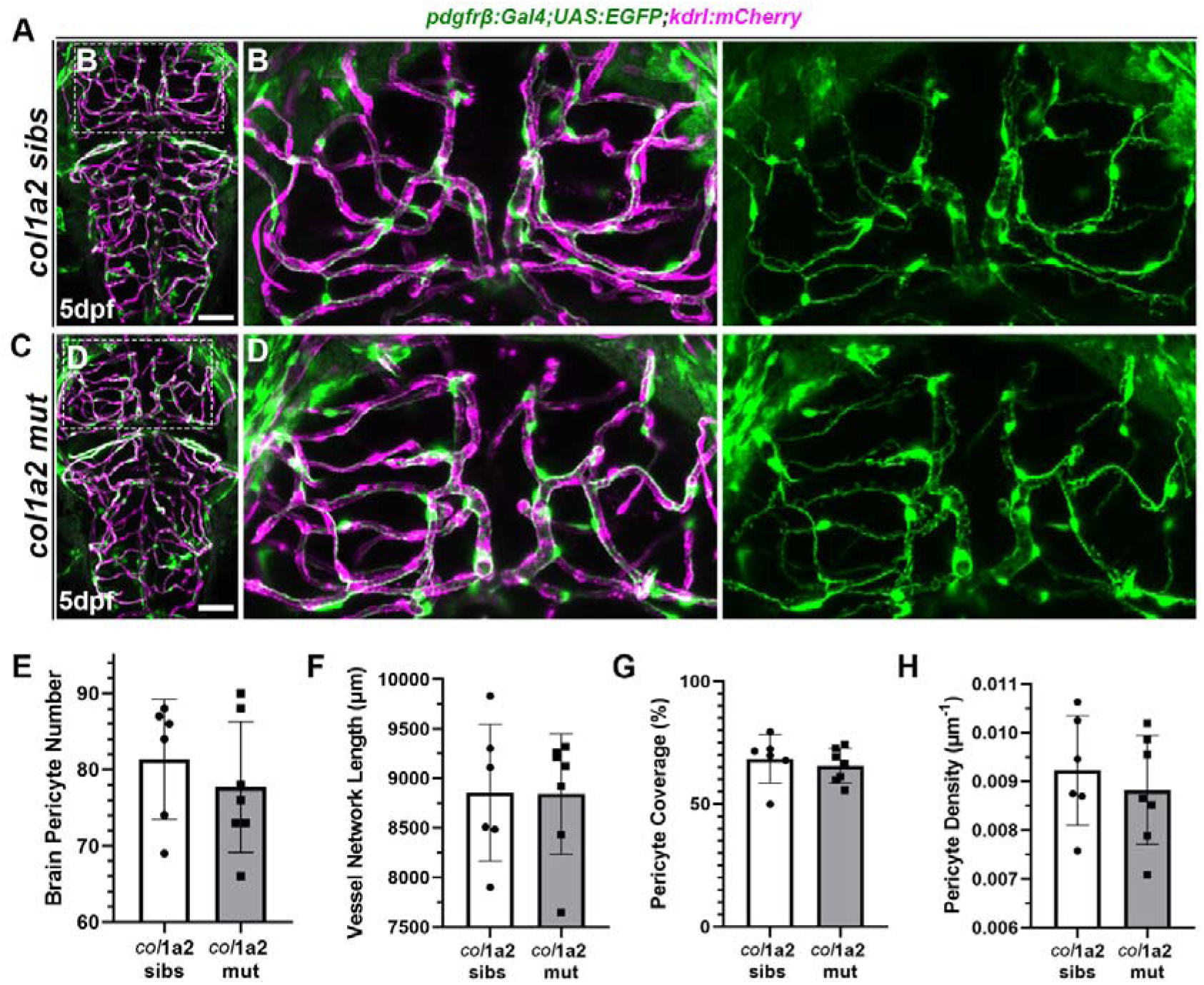
*col1a2^ca108^* mutants show no difference in pericyte number, coverage, and density. (A-D) Dorsal view of a 5dpf *col1a2^ca108^*(A-B) siblings and (C-D) homozygous mutants zebrafish larva on pericytes (pdgfrβ; green) and endothelial (*kdrl*; magenta) transgenic lines. (E-H) Quantification showing no significant difference in (E) pericyte number, (F) vessel network length, (G) pericyte coverage, and (H) pericyte density between *col1a2^ca108^* siblings and mutants (n: sibs= 6, mut= 7). Statistical analyses were performed using unpaired t-tests, with results presented as mean ± SD. Scale bars: 50µm.

**Fig. S8.**
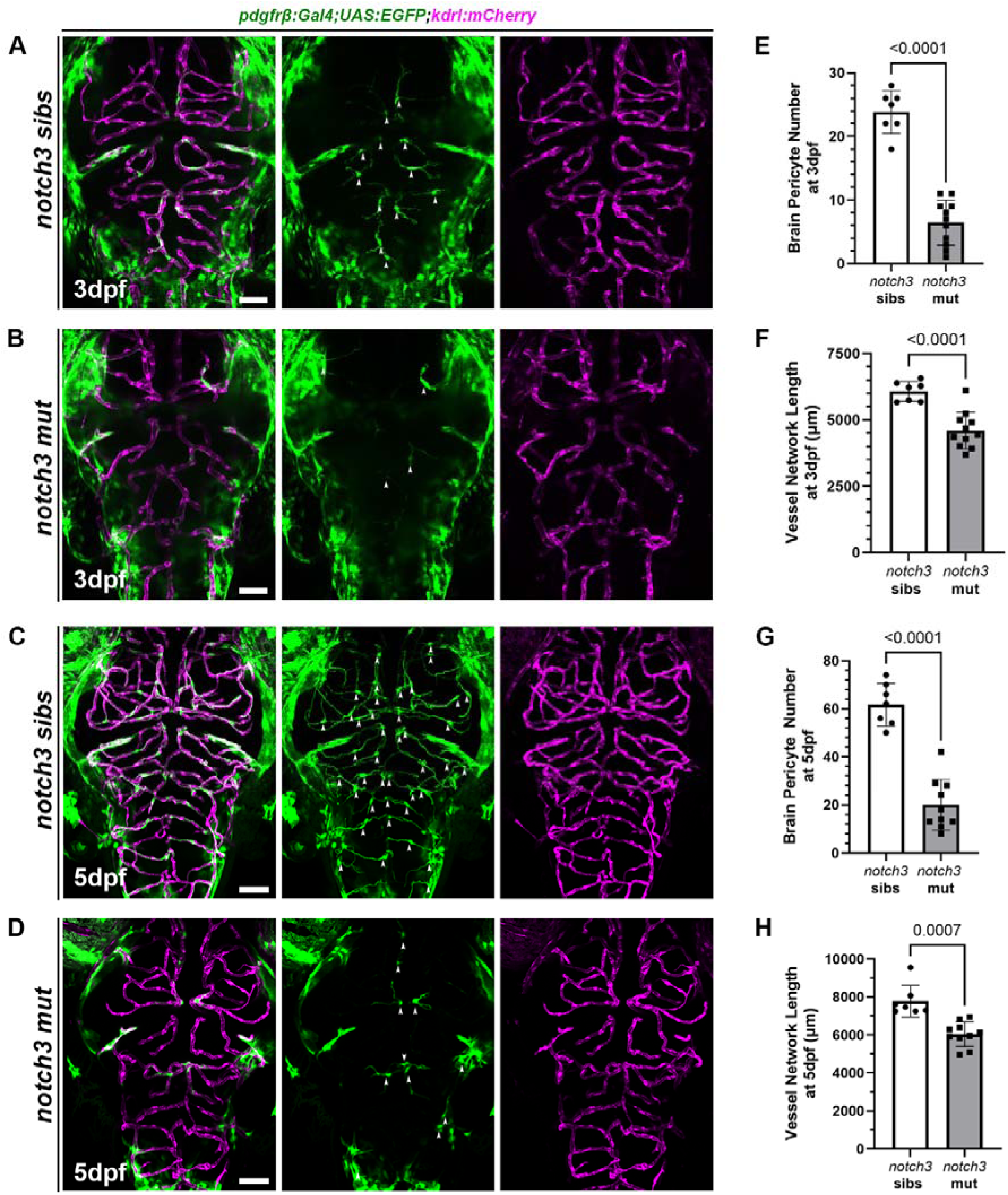
*notch3^fh322^* mutants have reduced pericyte numbers at 3 and 5dpf. (A-D) Confocal image of a 3dpf (A*) notch3^fh322^* siblings (B) homozygous mutant, and 5dpf (C*) notch3^fh322^* siblings (D) homozygous mutant zebrafish embryo on pericyte (green) and endothelial (magenta) transgenic lines. White arrows indicate pericytes in the image. (E-H) Quantification at 3dpf showing (E) reduced brain pericyte number and (F) vessel network length, and at 5dpf showing (G) reduced brain pericyte number and (H) vessel network length (3dpf n: sibs= 7, mut= 11, 5dpf n: sibs= 7, mut=9). Statistical analyses used unpaired t-tests, with results presented as mean ± SD. Scale bars: 50µm.

## REFERENCES

Abraham, S., Kogata, N., Fässler, R. and Adams, R. H. (2008). Integrin β1 Subunit Controls Mural Cell Adhesion, Spreading, and Blood Vessel Wall Stability. Circ. Res. 102, 562–570.

Ahuja, S., Adjekukor, C., Li, Q., Kocha, K. M., Rosin, N., Labit, E., Sinha, S., Narang, A., Long, Q., Biernaskie, J., et al. (2024). The development of brain pericytes requires expression of the transcription factor nkx3.1 in intermediate precursors. PLOS Biol. 22, e3002590.

Alunni, A., Krecsmarik, M., Bosco, A., Galant, S., Pan, L., Moens, C. B. and Bally-Cuif, L. (2013). Notch3 signaling gates cell cycle entry and limits neural stem cell amplification in the adult pallium. Development 140, 3335–3347.

Ando, K., Fukuhara, S., Izumi, N., Nakajima, H., Fukui, H., Kelsh, R. N. and Mochizuki, N. (2016). Clarification of mural cell coverage of vascular endothelial cells by live imaging of zebrafish. Development 143, 1328–1339.

Ando, K., Wang, W., Peng, D., Chiba, A., Lagendijk, A. K., Barske, L., Crump, J. G., Stainier, D. Y. R., Lendahl, U., Koltowska, K., et al. (2019). Peri-arterial specification of vascular mural cells from naïve mesenchyme requires Notch signaling. Development 146, dev165589.

Ando, K., Ishii, T. and Fukuhara, S. (2021). Zebrafish vascular mural cell biology: recent advances, development, and functions. Life 11, 1041.

Armulik, A., Genové, G., Mäe, M., Nisancioglu, M. H., Wallgard, E., Niaudet, C., He, L., Norlin, J., Lindblom, P. and Strittmatter, K. (2010). Pericytes regulate the blood–brain barrier. Nature 468, 557–561.

Armulik, A., Genové, G. and Betsholtz, C. (2011). Pericytes: Developmental, Physiological, and Pathological Perspectives, Problems, and Promises. Dev. Cell 21, 193–215.

Asakawa, K., Suster, M. L., Mizusawa, K., Nagayoshi, S., Kotani, T., Urasaki, A., Kishimoto, Y., Hibi, M. and Kawakami, K. (2008). Genetic dissection of neural circuits by Tol2 transposon-mediated Gal4 gene and enhancer trapping in zebrafish. Proc. Natl. Acad. Sci. 105, 1255–1260.

Bahrami, N. and Childs, S. J. (2020). Development of vascular regulation in the zebrafish embryo. Development 147, dev183061.

Benjamin, L. E., Hemo, I. and Keshet, E. (1998). A plasticity window for blood vessel remodelling is defined by pericyte coverage of the preformed endothelial network and is regulated by PDGF-B and VEGF. Development 125, 1591–1598.

Berthiaume, A.-A., Grant, R. I., McDowell, K. P., Underly, R. G., Hartmann, D. A., Levy, M., Bhat, N. R. and Shih, A. Y. (2018). Dynamic remodeling of pericytes in vivo maintains capillary coverage in the adult mouse brain. Cell Rep. 22, 8–16.

Berthiaume, A.-A., Schmid, F., Stamenkovic, S., Coelho-Santos, V., Nielson, C. D., Weber, B., Majesky, M. W. and Shih, A. Y. (2022). Pericyte remodeling is deficient in the aged brain and contributes to impaired capillary flow and structure. Nat. Commun. 13, 5912.

Brown, L. S., Foster, C. G., Courtney, J.-M., King, N. E., Howells, D. W. and Sutherland, B. A. (2019). Pericytes and Neurovascular Function in the Healthy and Diseased Brain. Front. Cell. Neurosci. Volume 13–2019,.

Chiaverina, G., di Blasio, L., Monica, V., Accardo, M., Palmiero, M., Peracino, B., Vara-Messler, M., Puliafito, A. and Primo, L. (2019). Dynamic Interplay between Pericytes and Endothelial Cells during Sprouting Angiogenesis. Cells 8, 1109.

Choi, J., Dong, L., Ahn, J., Dao, D., Hammerschmidt, M. and Chen, J.-N. (2007). FoxH1 negatively modulates flk1 gene expression and vascular formation in zebrafish. Dev. Biol. 304, 735–744.

Crouch, E. E., Bhaduri, A., Andrews, M. G., Cebrian-Silla, A., Diafos, L. N., Birrueta, J. O., Wedderburn-Pugh, K., Valenzuela, E. J., Bennett, N. K., Eze, U. C., et al. (2022). Ensembles of endothelial and mural cells promote angiogenesis in prenatal human brain. Cell 185, 3753–3769.e18.

Davison, J. M., Akitake, C. M., Goll, M. G., Rhee, J. M., Gosse, N., Baier, H., Halpern, M. E., Leach, S. D. and Parsons, M. J. (2007). Transactivation from Gal4-VP16 transgenic insertions for tissue-specific cell labeling and ablation in zebrafish. Dev. Biol. 304, 811–824.

DeSisto, J., O’Rourke, R., Jones, H. E., Pawlikowski, B., Malek, A. D., Bonney, S., Guimiot, F., Jones, K. L. and Siegenthaler, J. A. (2020). Single-Cell Transcriptomic Analyses of the Developing Meninges Reveal Meningeal Fibroblast Diversity and Function. Dev. Cell 54, 43–59.e4.

Dessalles, C. A., Babataheri, A. and Barakat, A. I. (2021). Pericyte mechanics and mechanobiology. J. Cell Sci. 134, jcs240226.

Dorsky, R. I., Sheldahl, L. C. and Moon, R. T. (2002). A Transgenic Lef1/β-Catenin-Dependent Reporter Is Expressed in Spatially Restricted Domains throughout Zebrafish Development. Dev. Biol. 241, 229–237.

Duan, L. and Yu, X. (2024). Fibroblasts: New players in the central nervous system? Fundam. Res. 4, 262–266.

Etchevers, H. C., Vincent, C., Douarin, N. M. L. and F. Couly, G. (2001). The cephalic neural crest provides pericytes and smooth muscle cells to all blood vessels of the face and forebrain. Development 128, 1059–1068.

Fleenor, B. S., Marshall, K. D., Durrant, J. R., Lesniewski, L. A. and Seals, D. R. (2010). Arterial stiffening with ageing is associated with transforming growth factor-β1-related changes in adventitial collagen: reversal by aerobic exercise. J. Physiol. 588, 3971–3982.

Gautam, J. and Yao, Y. (2018). Roles of Pericytes in Stroke Pathogenesis. Cell Transplant. 27, 1798–1808.

Goss, G., Rognoni, E., Salameti, V. and Watt, F. M. (2021). Distinct fibroblast lineages give rise to NG2+pericyte populations in mouse skin development and repair. Front. Cell Dev. Biol. 9, 675080.

Graff, M. F. E., Heeg, E. E. and Childs, S. J. (2025). Progressive mural cell deficiencies across the lifespan in a foxf2 model of Cerebral Small Vessel Disease.

Grant, R. I., Hartmann, D. A., Underly, R. G., Berthiaume, A.-A., Bhat, N. R. and Shih, A. Y. (2019). Organizational hierarchy and structural diversity of microvascular pericytes in adult mouse cortex. J. Cereb. Blood Flow Metab. 39, 411–425.

Greisenegger, E. K., Llufriu, S., Chamorro, A., Cervera, A., Jimenez-Escrig, A., Rappersberger, K., Marik, W., Greisenegger, S., Stögmann, E. and Kopp, T. (2021). A NOTCH3 homozygous nonsense mutation in familial Sneddon syndrome with pediatric stroke. J. Neurol. 268, 810–816.

Hellström, M., Kalén, M., Lindahl, P., Abramsson, A. and Betsholtz, C. (1999). Role of PDGF-B and PDGFR-β in recruitment of vascular smooth muscle cells and pericytes during embryonic blood vessel formation in the mouse. Development 126, 3047–3055.

Herbomel, P., Thisse, B. and Thisse, C. (2001). Zebrafish Early Macrophages Colonize Cephalic Mesenchyme and Developing Brain, Retina, and Epidermis through a M-CSF Receptor-Dependent Invasive Process. Dev. Biol. 238, 274–288.

Hielscher, A., Ellis, K., Qiu, C., Porterfield, J. and Gerecht, S. (2016). Fibronectin Deposition Participates in Extracellular Matrix Assembly and Vascular Morphogenesis. PLOS ONE 11, e0147600.

Jiang, X., Iseki, S., Maxson, R. E., Sucov, H. M. and Morriss-Kay, G. M. (2002). Tissue Origins and Interactions in the Mammalian Skull Vault. Dev. Biol. 241, 106–116.

Jones, H. E., Coelho-Santos, V., Bonney, S. K., Abrams, K. A., Shih, A. Y. and Siegenthaler, J. A. (2023). Meningeal origins and dynamics of perivascular fibroblast development on the mouse cerebral vasculature. Development 150, dev201805.

Jones, H. E., Abrams, K. A., Kim, S., Fantauzzo, K. A. and Siegenthaler, J. A. (2026). PDGFRα is required for postnatal cerebral perivascular fibroblast development. Dev. Biol. 530, 1–11.

Kelly, K. K., MacPherson, A. M., Grewal, H., Strnad, F., Jones, J. W., Yu, J., Pierzchalski, K., Kane, M. A., Herson, P. S. and Siegenthaler, J. A. (2016). Col1a1+ perivascular cells in the brain are a source of retinoic acid following stroke. BMC Neurosci. 17, 49.

Lee, U., Zhang, Y., Zhu, Y., Luo, A. C., Gong, L., Tremmel, D. M., Kim, Y., Villarreal, V. S., Wang, X., Lin, R.-Z., et al. (2024). Robust differentiation of human pluripotent stem cells into mural progenitor cells via transient activation of NKX3.1. Nat. Commun. 15, 8392.

Lin, P. K. and Davis, G. E. (2023). Extracellular Matrix Remodeling in Vascular Disease: Defining Its Regulators and Pathological Influence. Arterioscler. Thromb. Vasc. Biol. 43, 1599–1616.

Lindahl, P., Johansson, B. R., Levéen, P. and Betsholtz, C. (1997). Pericyte Loss and Microaneurysm Formation in PDGF-B-Deficient Mice. Science 277, 242–245.

Lu, S., Jolly, A. J., Strand, K. A., Dubner, A. M., Mutryn, M. F., Moulton, K. S., Nemenoff, R. A., Majesky, M. W. and Weiser-Evans, M. C. M. (2020). Smooth muscle–derived progenitor cell myofibroblast differentiation through KLF4 downregulation promotes arterial remodeling and fibrosis. JCI Insight 5,.

Ma, R. C., Jacobs, C. T., Sharma, P., Kocha, K. M. and Huang, P. (2018). Stereotypic generation of axial tenocytes from bipartite sclerotome domains in zebrafish. PLOS Genet. 14, e1007775.

McGarry, S. D., Adjekukor, C., Ahuja, S., Greysson-Wong, J., Vien, I., Rinker, K. D. and Childs, S. J. (2024). Vessel Metrics: A software tool for automated analysis of vascular structure in confocal imaging. Microvasc. Res. 151, 104610.

O’Brown, N. M., Megason, S. G. and Gu, C. (2019). Suppression of transcytosis regulates zebrafish blood-brain barrier function. eLife 8, e47326.

Payne, L. B., Darden, J., Suarez-Martinez, A. D., Zhao, H., Hendricks, A., Hartland, C., Chong, D., Kushner, E. J., Murfee, W. L. and Chappell, J. C. (2021). Pericyte migration and proliferation are tightly synchronized to endothelial cell sprouting dynamics. Integr. Biol. 13, 31–43.

Pippucci, T., Maresca, A., Magini, P., Cenacchi, G., Donadio, V., Palombo, F., Papa, V., Incensi, A., Gasparre, G., Valentino, M. L., et al. (2015). Homozygous NOTCH3 null mutation and impaired NOTCH3 signaling in recessive early-onset arteriopathy and cavitating leukoencephalopathy. EMBO Mol. Med. 7, 848–858.

Posokhova, E., Shukla, A., Seaman, S., Volate, S., Hilton, M. B., Wu, B., Morris, H., Swing, D. A., Zhou, M. and Zudaire, E. (2015). GPR124 functions as a WNT7-specific coactivator of canonical β-catenin signaling. Cell Rep. 10, 123–130.

Pouget, C., Pottin, K. and Jaffredo, T. (2008). Sclerotomal origin of vascular smooth muscle cells and pericytes in the embryo. Dev. Biol. 315, 437–447.

Proulx, K., Lu, A. and Sumanas, S. (2010). Cranial vasculature in zebrafish forms by angioblast cluster-derived angiogenesis. Dev. Biol. 348, 34–46.

Rajan, A. M., Ma, R. C., Kocha, K. M., Zhang, D. J. and Huang, P. (2020). Dual function of perivascular fibroblasts in vascular stabilization in zebrafish. PLoS Genet. 16, e1008800.

Rajan, A. M., Rosin, N. L., Labit, E., Biernaskie, J., Liao, S. and Huang, P. (2023). Single-cell analysis reveals distinct fibroblast plasticity during tenocyte regeneration in zebrafish. Sci. Adv. 9, eadi5771.

Shen, J., Ju, D., Wu, S., Zhao, J., Pham, L., Ponce, A., Yang, M., Li, H. J., Zhang, K., Yang, Z., et al. (2024). SM22α deficiency: promoting vascular fibrosis via SRF-SMAD3-mediated activation of Col1a2 transcription following arterial injury.

Shih, Y.-H., Portman, D., Idrizi, F., Grosse, A. and Lawson, N. D. (2021). Integrated molecular analysis identifies a conserved pericyte gene signature in zebrafish. Development 148, dev200189.

Speir, M. L., Bhaduri, A., Markov, N. S., Moreno, P., Nowakowski, T. J., Papatheodorou, I., Pollen, A. A., Raney, B. J., Seninge, L., Kent, W. J., et al. (2021). UCSC Cell Browser: visualize your single-cell data. Bioinformatics 37, 4578–4580.

Stellingwerff, M. D.⍰; N., Corinne; Helman, Guy; Roosendaal, Stefan D.⍰;. Benko, William S.⍰;. Pizzino, Amy; Bugiani, Marianna; Vanderver, Adeline; Simons, Cas; van der Knaap, Marjo S. (2022). Early-Onset Vascular Leukoencephalopathy Caused by Bi-Allelic NOTCH3 Variants. Neuropediatrics 53, 115–121.

Sun, Z., Gao, C., Gao, D., Sun, R., Li, W., Wang, F., Wang, Y., Cao, H., Zhou, G., Zhang, J., et al. (2021). Reduction in pericyte coverage leads to blood–brain barrier dysfunction via endothelial transcytosis following chronic cerebral hypoperfusion. Fluids Barriers CNS 18, 21.

Sur, A., Wang, Y., Capar, P., Margolin, G., Prochaska, M. K. and Farrell, J. A. (2023). Single-cell analysis of shared signatures and transcriptional diversity during zebrafish development. Dev. Cell 58, 3028–3047.e12.

Tarallo, S., Beltramo, E., Berrone, E. and Porta, M. (2012). Human pericyte–endothelial cell interactions in co-culture models mimicking the diabetic retinal microvascular environment. Acta Diabetol. 49, 141–151.

Vanhollebeke, B., Stone, O. A., Bostaille, N., Cho, C., Zhou, Y., Maquet, E., Gauquier, A., Cabochette, P., Fukuhara, S., Mochizuki, N., et al. (2015). Tip cell-specific requirement for an atypical Gpr124- and Reck-dependent Wnt/β-catenin pathway during brain angiogenesis. eLife 4, e06489.

Vanlandewijck, M., He, L., Mäe, M. A., Andrae, J., Ando, K., Del Gaudio, F., Nahar, K., Lebouvier, T., Laviña, B. and Gouveia, L. (2018). A molecular atlas of cell types and zonation in the brain vasculature. Nature 554, 475–480.

Wang, Y., Pan, L., Moens, C. B. and Appel, B. (2014). Notch3 establishes brain vascular integrity by regulating pericyte number. Development 141, 307–317.

Whitesell, T. R., Kennedy, R. M., Carter, A. D., Rollins, E.-L., Georgijevic, S., Santoro, M. M. and Childs, S. J. (2014). An α-Smooth Muscle Actin (acta2/αsma) Zebrafish Transgenic Line Marking Vascular Mural Cells and Visceral Smooth Muscle Cells. PLOS ONE 9, e90590.

Whitesell, T. R., Chrystal, P. W., Ryu, J.-R., Munsie, N., Grosse, A., French, C. R., Workentine, M. L., Li, R., Zhu, L. J., Waskiewicz, A., et al. (2019). foxc1 is required for embryonic head vascular smooth muscle differentiation in zebrafish. Dev. Biol. 453, 34–47.

Yang, A. C., Vest, R. T., Kern, F., Lee, D. P., Agam, M., Maat, C. A., Losada, P. M., Chen, M. B., Schaum, N., Khoury, N., et al. (2022). A human brain vascular atlas reveals diverse mediators of Alzheimer’s risk. Nature 603, 885–892.

Zhou, Y. and Nathans, J. (2014). Gpr124 Controls CNS Angiogenesis and Blood-Brain Barrier Integrity by Promoting Ligand-Specific Canonical Wnt Signaling. Dev. Cell 31, 248–256.

Zi, H., Peng, X., Cao, J., Xie, T., Liu, T., Li, H., Bu, J., Du, J. and Li, J. (2024). Piezo1-dependent regulation of pericyte proliferation by blood flow during brain vascular development. Cell Rep. 43,.

