## Supplemental tables for "Dissecting embryonic origins of brain pericytes, the role of angiogenesis, and fibroblasts’ contributions"

Table S1 Morpholino sequences

| **Molecule** | **Sequence** | **Identifier** | **Dose (ng/embryo)** | **Reference** |
| --- | --- | --- | --- | --- |
| Gpr124 MO | ACTGATATTGATTTAACTCACCACA | GENE TOOLS, LLC | 0.8 | (Vanhollebeke et al., 2015) |

Table S2 Drug list and their working concentration

| **Drug** | **Company, Cat No.** | **Working Concentration** | **Target Pathway** |
| --- | --- | --- | --- |
| DMH4 | SIGMA, D8696 | 10µM | Inhibits Vegfr2 |
| SU5416 | SIGMA, S8442 | 1.5µM | Inhibits Vegfr1/2 |
| Metronidazole | SIGMA, M3761 | 5mM | Cause cell death in nitroreductase (NTR) positive cells. |
| DMSO | SIGMA, D8418 | Equivalent dilution of drug |  |

Table S3 Antibodies and dilutions

| **Antibody** | **Company, Cat No.** | **Host Specie** | **Dilution** |
| --- | --- | --- | --- |
| GFP | Clontech, 632381 | Mouse | 1:500 |
| Phosphohistone H3 | Sigma-Aldrich, 06-570 | Rabbit | 1:250 |
| Alexa Fluor 488 | Invitrogen, A-21202 | Donkey α-mouse | 1:500 |
| Alexa Fluor 647 | Invitrogen, A-21244 | Donkey α-Rabbit | 1:500 |

Table S4 HCR molecular instruments probes

| **Gene** | **Transcript Ref No.** | **Amplifier** | **Flourophore** | **Dilution** |
| --- | --- | --- | --- | --- |
| *col1a2* | ZDB-GENE-030131-8415 | X2 | Alexa 546 | 2:100 |
| *col5a1* | ZDB-GENE-041105-6 | X1 | Alexa 488 | 2.5:100 |
| *kdrl* | ZDB-GENE-000705-1 | B2 | Alexa 647 | 2:100 |
| *pdgfrb* | ZDB-GENE-030805-2 | B4 | Alexa 488 | 2:100 |
